# Strigolactone effects on *Sorghum bicolor* ecophysiology and symbioses

**DOI:** 10.1101/2025.08.07.669140

**Authors:** Chloee M. McLaughlin, Margarita Takou, Joel Masanga, Erica H. Lawrence-Paul, Evelyn J. Abraham, Melanie Perryman, Anna Calabritto, Amandeep Cheema, Yuxing Xu, Baloua Nebie, Steven Runo, Joshua J. Kellogg, Roberta Croce, Daniel P. Schachtman, Huirong Gao, Ruairidh J. H. Sawers, Jesse R. Lasky

## Abstract

Strigolactones are ecologically, developmentally, and physiologically important hormones, but much remains unknown about their evolution and role in non-model species. Sorghum is a globally important C_4_ cereal and exhibits natural variation in root-exuded strigolactones. Differences in sorghum strigolactone stereochemistry are associated with resistance to parasitic plants, but with evidence for potential trade-offs. We studied sorghum mutants of loci in the strigolactone biosynthetic pathway, C*AROTENOID CLEAVAGE DIOXYGENASE 8* (Sb*CCD8b*) and *LOW GERMINATION STIMULANT 1* (*LGS1*), previously shown to be *Striga* resistant by stimulating little germination of the parasite. Sb*CCD8b* CRISPR-Cas9 deletions changed the accumulation of low abundance metabolites, reduced net carbon assimilation rate, altered root architecture and anatomy, and diminished the establishment and benefit of mycorrhizal symbionts. For *Striga-*resistant *LGS1* CRISPR-Cas9 deletions, differentially expressed genes were enriched with promoter motifs for stress response and growth pathways, net carbon assimilation rate was reduced, and the colonization of mycorrhizal symbionts was delayed. We additionally restored functional *LGS1* into the RTx430 genetic background, which normally has the *lgs1*-2 natural deletion allele. While root exudates from *LGS1* insertion mutants rescued *Striga* susceptibility, we did not see consistent rescue of other traits impacted in *LGS1* loss-of-function mutants. We hypothesize that epistasis with a neighboring strigolactone synthesis gene, which is rarely lost without concomitant loss of *LGS1,* may alter the phenotypic effects of *LGS1* variation. Our study gives context to potential trade-offs associated with host resistance to parasitic plants and, more broadly, builds on the contribution of strigolactones in shaping sorghum physiological processes, growth, and development.

## Introduction

Strigolactones, originally named for their role as germination stimulants to the parasitic plant *Striga* spp. (Cook et al., 1966) are well established as an important class of hormones for plant biology and ecological interactions (Saeed et al., 2017; Wu et al., 2022). Strigolactones play key roles in various aspects of plant growth, physiology, and symbiotic interactions, including the regulation of shoot architecture, root development, light harvesting complexes, and associations with mycorrhizal fungi (Akiyama et al., 2005; Gomez-Roldan et al., 2008; Mayzlish-Gati et al., 2010; Ruyter-Spira et al., 2011). As signaling molecules, strigolactones enable plants to regulate their growth and development in response to the external environment (Barbier et al., 2023). To date, more than 30 strigolactones have been characterized which differ in their structure (Yoneyama and Brewer, 2021), though much of the functional variation associated with this structural diversity remains to be elucidated.

Strigolactones are synthesized from carotenoids which undergo catalytic reactions by several enzymes. Their biosynthesis begins with the isomerization of β-carotene from a trans to a cis configuration by β-carotene isomerase (*DWARF27*, *D27*). 9-cis-β-carotene is then cleaved by two carotenoid cleavage dioxygenases, *CAROTENOID CLEAVAGE DIOXYGENASE 7* (*CCD7*) and *CAROTENOID CLEAVAGE DIOXYGENASE 8* (*CCD8*), resulting in the formation of carlactone (Alder et al., 2012; Seto et al., 2014). Mutants of *CCD7* and *CCD8* have undetectable or extremely reduced exudation of strigolactones (tomato, Sl*CCD8* (Kohlen et al., 2012); sorghum, Sb*CCD8b* (Hao et al., 2023); rice, Os*CCD7* (Butt et al., 2018)), demonstrating that *CCD*s function as critical enzymes for the biosynthesis of strigolactones.

Mutants of *CCD7* and *CCD8* homologs are not only deficient in strigolactone exudation, but also have altered shoot tillering and branching (*Arabidopsis* (Sorefan et al., 2003); rice (Arite et al., 2007); tobacco (Gao et al., 2018); tomato (Vogel et al., 2010); maize (Guan et al., 2012); sorghum (Hao et al., 2023)) and altered root system architecture (rice (Arite et al., 2012); tomato (Kohlen et al., 2012); *Arabidopsis* and pea (Rasmussen et al., 2012); maize (Guan et al., 2012); sorghum (Hao et al., 2023)). More recently, strigolactone signaling has been linked to variation in root anatomy. For example, (Zhao et al., 2025) showed that strigolactones negatively regulate cambium-derived xylem vessel formation in *Arabidopsis*, while (Shrestha et al., 2026) demonstrated strigolactone-deficient Os*PSY2* mutants in rice to have impaired aerenchyma formation that can be rescued by exogenous strigolactone application. Beyond modulating plant architecture and morphology, strigolactones also play a role in shaping physiological traits and have been linked to differences in stomatal development and behavior (Van Ha et al., 2014), chlorophyll content and the efficiency of light capture through transcriptional regulation (Mayzlish-Gati et al., 2010). Transcriptome analysis of strigolactone deficient tomato mutants (Sl*ORT1*) revealed decreased expression of genes involved in light harvesting which can be rescued with exogenous treatment of the synthetic strigolactone analog GR24 (Mayzlish-Gati et al., 2010), suggesting that strigolactones are positive regulators of light harvesting processes. Altogether, the number of plant phenotypes regulated by or associated with differences in strigolactone profiles establishes their vital role in plant response to the environment and downstream impacts on numerous complex traits.

Along with other plant hormones, strigolactones play a key role in signaling pathways that integrate nutrient and environmental cues to form beneficial symbioses. For example, plants have evolved the phosphate starvation response (PSR) which reprograms plant processes to acquire more phosphorus from the environment and to remobilize phosphorus under nutrient-deprived conditions (Zhang et al., 2014; Chien et al., 2018). PSR results in an increase of synthesis and exudation of strigolactones (Wang et al., 2021; Haider et al., 2023). This increase in strigolactones provides two crucial functions: first to serve as endogenous hormones that control plant development and stress response, and secondly, to serve as exogenous signaling molecules and promote symbiosis with beneficial arbuscular mycorrhizal fungi (AMF). AMF are obligate heterotrophs that interact with the roots of more than 80 percent of land plants, including crop plants, and provide difficult to access nutrients in return for photosynthetically fixed carbon and lipids (Brundrett and Tedersoo, 2018). AMF colonizes the host root cortex before differentiating into highly branched structures called arbuscules, the main site of nutrient exchange between the fungal and plant symbiotic partners. Strigolactones play a key role in the initiation of the symbiosis by inducing AMF hyphal branching, enhancing the possibility of contact between AMF and a host root (Giovannetti et al., 1993; Giovannetti et al., 1994; Kodama et al., 2022).

Host plant exudation of strigolactones to recruit AMF symbionts has been hijacked by parasitic plant members of the Orobanchaceae family. Obligate root parasites in this family germinate after sensing host root exuded strigolactones (Yoneyama et al., 2010) and parasites in this family generally use host strigolactones as chemoattractants (Ogawa et al., 2022). Of particular interest is the obligate root parasite *Striga hermonthica,* which accounts for billions of dollars’ worth of annual cereal crop damage in tropical Africa and is considered one of the greatest constraints to African food security (Spallek et al., 2013). One of the best characterized host resistance mechanisms is low germination of *S. hermonthica* by the host’s root exudates (Gurney et al., 2003; Ruyter-Spira et al., 2011; Mallu et al., 2021). In sorghum, naturally occurring deletion mutants of the *LOW GERMINATION STIMULANT 1* (*LGS1*) quantitative trait locus (QTL) confer shifts in strigolactone stereochemistry, where orobanchol-type rather than 5-deoxystrigol (5DS) becomes the more abundant strigolactone in root exudates (Gobena et al., 2017). The gene causing the low germination stimulation phenotype (*Sobic.005213600, LGS1*) encodes a sulfotransferase that functions downstream of CYP711A (Sb*MAX1a*) converting 18-hydroxy-carlactonoic acid (18-hydroxy-CLA) into 5DS and 4-deoxyorobanchol (4DO) simultaneously (Wu and Li, 2021; Yoda et al., 2021). In addition to altering the strigolactone composition, loss-of-function at the *LGS1* gene may reduce the quantity of total strigolactone exudation (Bellis et al., 2020). Directly upstream of the *LGS1* gene is *Sobic.005G213500* (*Sb3500*), a 2-oxoglutarate-dependent dioxygenase involved in stereoselective strigolactone production and increases the quantity of 5DS relative to 4DO when expressed with *LGS1* (Yoda et al., 2023). Previously described naturally occurring deletion variants at the QTL either lack both *LGS1* and *Sb3500* (e.g. *lgs1-*1, *lgs1*-2), or retain *Sb3500* with absence of *LGS1* (e.g. *lgs1*-3) (Gobena et al., 2017). Thus, the genomic region where *LGS1* resides may function as a multi-gene module where epistasis influences strigolactone profiles, though much of the work to date on *LGS1*-based resistance in sorghum has focused on the gene encoding the sulfotransferase (*LGS1*, *Sobic.005213600*).

Despite the potential for loss-of-function in the *LGS1* gene to increase sorghum’s fitness in landscapes with parasite pressure, the alterations to strigolactone biosynthesis may impair endogenous processes in sorghum. Supporting this hypothesis, Bellis et al., (2020) found evidence of potential trade-offs: natural loss-of-function *lgs1* alleles are maintained at a modest frequency in sorghum landraces within *S. hermonthica*-prone regions (up to ∼30% in northern Nigeria) and are at very low frequency in *S. hermonthica-*free landscapes. Also, CRISPR-Cas9 deletions of *LGS1* altered expression of >2,000 genes compared to the wild type, and there was a significant downregulation of light harvesting genes in shoots (Bellis et al., 2020). Given the broad roles strigolactones play in plant processes, we hypothesized that sorghum with loss-of-function at *LGS1* is subject to trade-offs through pleiotropic impacts.

In the present study, we explore the impact of mutations at two strigolactone biosynthesis genes, Sb*CCD8b* and *LGS1,* on sorghum physiology, growth, and biotic interactions with parasitic plants and arbuscular mycorrhizal fungi to ask the following questions: (Q1) How does variation at strigolactone biosynthesis genes in sorghum impact biotic interactions with *Striga hermonthica* and arbuscular mycorrhizae fungi? (Q2) What regulatory pathways may be altered in mutants of strigolactone biosynthesis genes and are phenotypic changes consistent with altered pathways? (Q3) How does variation at strigolactone biosynthesis genes in sorghum affect sorghum performance and physiology? (Q4) Are strigolactone impacts consistent across diverse genetic backgrounds?

## Materials & Methods

### Generation of mutants

We studied loss-of-function in the genes Sb*CCD8* and *LGS1* using gene-edited *S. bicolor* of two different genetic backgrounds (Table 1). CRISPR-Cas9 edited lines of Sb*CCD8b* in the ‘RTx430’ background were included in our study to represent a strigolactone deficient line (RTx430 *ccd8b*, see greater detail in Hao et al., 2023). Native sorghum ecotypes have natural variation at the *LGS1* locus, impacting its functionality, and we included genetic two backgrounds, Macia and RTx430, to determine if *LGS1* effects are consistent. Functional *LGS1* was knocked out in the normally *LGS1*-functional background Macia (PI 565121) using CRISPR-Cas9 to produce independent genome-edited lines (Macia *lgs1*-d, see greater detail for generation of S_2_ plants in Bellis et al. 2020). Unless otherwise indicated, we used a more advanced version of the material described in Bellis et al. (2020) which was backcrossed to the recurrent background Macia and selfed for an additional generation (BC_0_F_2_) in this study (Figure S1, Supplementary Methods).

**Table 1.** Description of plant material used in this study. All gene models are reported from the BTx623v5.1 reference genome.

| Genetic Background | Genotype | Description |
| --- | --- | --- |
| RTx430 | RTx430 *ccd8b* | CRISPR/Cas9-edited deletion mutant of the *SbCCD8b* (*Sobic.003G293600*) gene described in Hao et al., (2023). |
|  | RTx430 WT (*lgs1-2*) | Wild type with the *lgs1-2* natural loss-of-function allele. The *lgs1-2* deletion allele spans the three gene models *Sobic.005G213400*, *Sb3500* (*Sobic.005G213500*), and *LGS1* (*Sobic.005G213600*). |
|  | RTx430 WT-L (*lgs1-2*) | Insertion transformation event null segregant. |
|  | RTx430 *LGS1* | Mutants with functional *LGS1* (*Sobic.005G213600*) introduced via *Agrobacterium* random transformation, described in this study. |
| Macia | Macia WT | Wild type with intact *LGS1* (*Sobic.005G213600*). |
|  | Macia WT-L | Deletion transformation event null segregant. |
|  | Macia *lgs1-d* | CRISPR/Cas9-edited deletion mutant of the gene *Sobic.005G213600* (*LGS1*). Material grown in this study includes the S<sub>2</sub> generation (described in Bellis et al., (2020)) and a more advanced generation backcrossed to the Macia wild type to generate BC<sub>0</sub>F<sub>2</sub> seed (Figure S1). |

We additionally sought to test the effect of inserting functional *LGS1* alone back into a normally non-functional background. RTx430 (PI 655996) is an inbred background, developed at Texas A&M using a partially converted Ethiopian sorghum variety (Miller, 1984). The RTx430 background carries the *lgs1-2* allele, which contains a large deletion that spans three predicted genes, including the sulfotransferase *LGS1* and the 2-oxoglutarate-dependent dioxygenase *Sb3500* (Gobena et al., 2017). *LGS1* was inserted via transformation with *Agrobacterium* into the RTx430 genetic background, to produce two independent mutants with restored functionality of the locus (RTx430 *LGS1*) (Supplementary Methods). For both *LGS1* insertion (RTx430) and deletion events (Macia), null event mutants were also produced and included in our analysis as “wild-type like” (WT-L) comparisons. See Table 1 for descriptions of the plant material used in this study.

### Evaluation of allelic diversity in natural sorghum populations

Because Yoda et al. (2023) found the effect of presence-absence of *LGS1* (*Sobic.005213600*) and *Sb3500* (*Sobic.005213500*) had interactive effects on strigolactones when transiently expressed in *Nicotiana benthamiana*, we evaluated patterns of presence-absence variation and linkage of these genes in sorghum landraces (Table 2)(Morris et al., 2026). Gene presence was determined using a local exon-level *de novo* reassembly pipeline, in which sequencing reads were assembled without mapping to a primary reference. Exon contigs including short flanking regions were stitched according to the BTx623v5.1 (Morris et al., 2026) exon order to reconstruct pseudo-mRNA sequences. Genes with reassembled proteins showing ≥90% identity and ≥90% coverage relative to BTx623v5.1 were classified as present. Strigolactone profiles reported in Table 2 were not directly measured, but inferred based on published functional annotations of *LGS1* and *Sb3500* and their roles in strigolactone biosynthesis (Gobena et al., 2017; Yoda et al., 2021; Yoda et al., 2023).

**Table 2.**
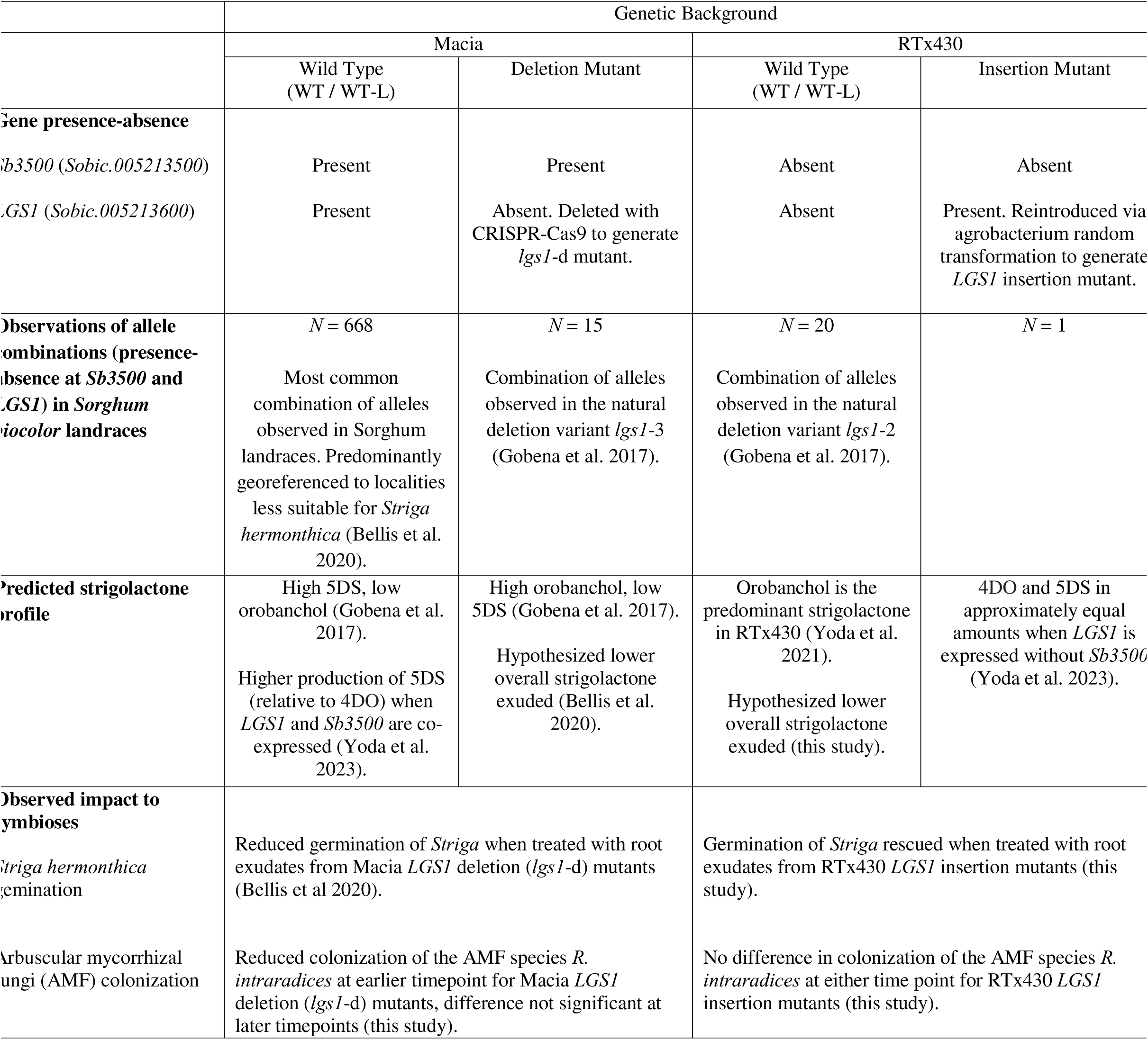
For material of the *LGS1* gene used in this study, summary of gene presence-absence at *Sb3500* and *LGS1*, observations of the allele combinations in landraces, predicted strigolactone profiles, and observed effects of synthetic mutants on symbiotic interactions.

### Striga hermonthica *germination assay*

Root exudates of the mutants with functional *LGS1* inserted, “wild type-like” (WT-L) null segregants, and the WT in the RTx430 background were collected to test their *S. hermonthica* germination activity. Plants were grown in a greenhouse in University Park, PA (16-hour photoperiod; day temperature 28°C; night temperature 22°C). Sorghum seeds were surface sterilized in 1.2% sodium hypochlorite for 10 minutes, rinsed with sterile deionized water (diH_2_O) and germinated in autoclaved vermiculite. One-week old seedlings were transferred to a hydroponics system comprising 50 mL falcon tubes filled with half-strength modified Hoagland solution. Each seedling was suspended in the solution by a bottomless 1.5 mL centrifuge tube held at the center of the 50 mL falcon tube cap (Wang et al., 2022). Plants were grown in this medium for seven days and tubes were manually agitated every other day to improve airflow. The medium was discarded, then tubes and seedling roots rinsed with diH_2_O. Tubes were refilled with half-strength Hoagland’s solution without phosphate, and seedlings grown for an additional 7 days. Root exudates were collected in 50 mL diH_2_O over a 48-hr period under darkness. Each seedling’s root volume was recorded by water displacement in a graduated cylinder, and exudate concentrations standardized to the lowest recorded root volume prior to use for induction of *S. hermonthica* germination.

Germination assays used *S. hermonthica* seed collected in 2016 from a sorghum field in Kibos, Kenya (0°40′S; 34°49′E) and in 2018 from a sorghum field in Siby, Mali (12°23′N; 8°20′W). Since *S. hermonthica* requires conditioning to be responsive to germination stimulants (Matusova et al., 2004), parasite seeds were surface sterilized in 1.2% sodium hypochlorite with a drop of Tween 20 for 20 mins, rinsed thoroughly with sterile diH_2_O, and incubated at 30°C in the dark for 10 days. After conditioning, the *S. hermonthica* seed-diH_2_O mixture was homogenized before pipetting 100 μL (approximately 100 seeds) into a single well of a 12-well microtiter plate (CytoOne). Excess water was aspirated out of the wells before adding 3 mL of sorghum root exudates. 15 biological replicates of each sorghum genotype, and 2 technical reps (wells per biological rep) were tested for germinability of each *S. hermonthica* population. A similar volume of 0.1ppm GR24 (synthetic strigolactone standard) and diH_2_O were used as positive and negative controls, respectively. *S. hermonthica* germination, characterized by emergence of a radicle, was checked after incubating the plates for 48 hours at 30°C in the dark, and quantified by calculating the percentage of germinated seeds in each well. Germination percentage of *S. hermonthica* in Figure 1 is plotted with a broken axis using functions described in R/ggbreak (Xu et al., 2021).

**Figure 1.**
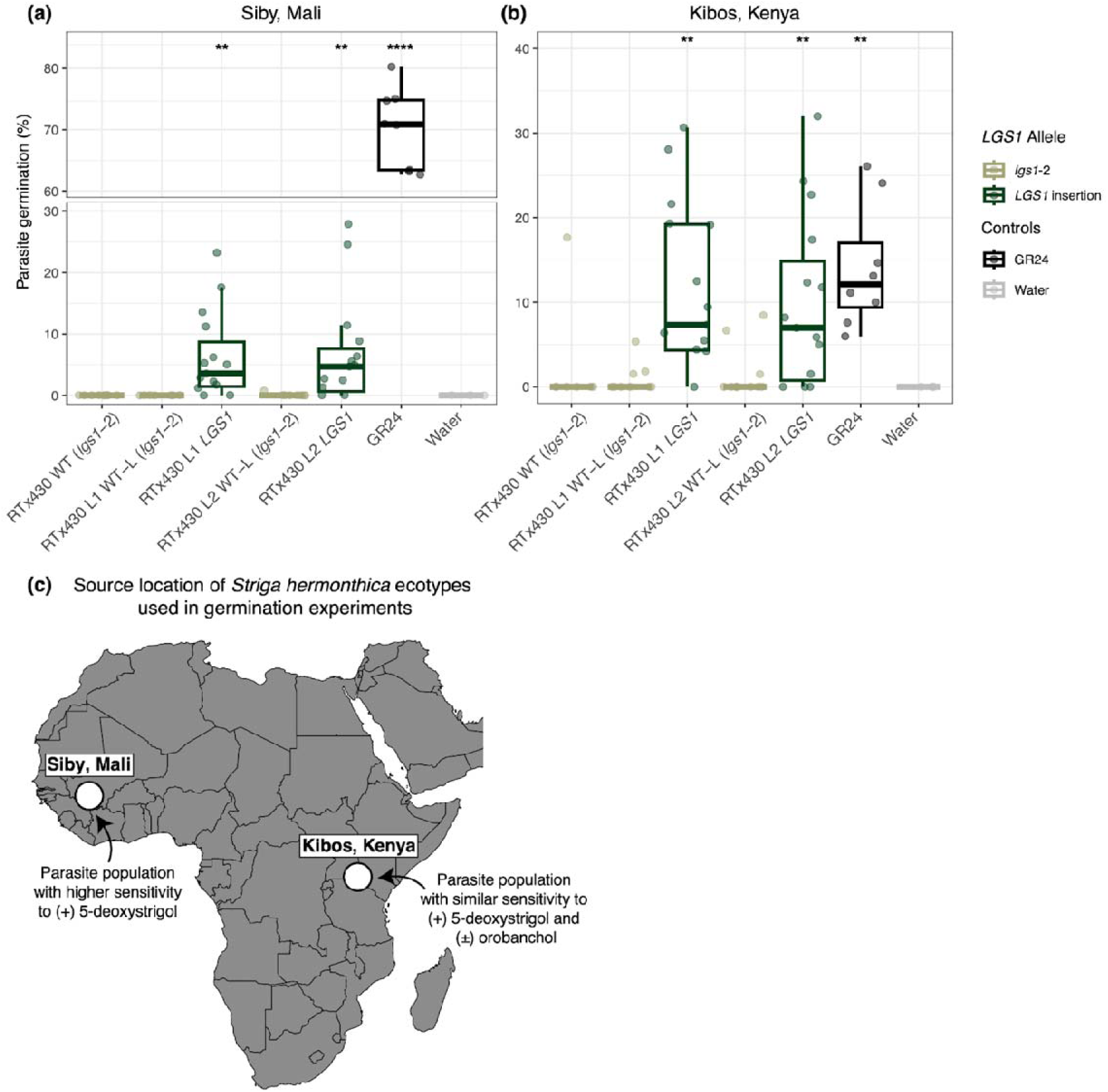
Restoring functional *LGS1* in the RTx430 background rescues *Striga hermonthica* germination. Percent *S. hermonthica* germination of ecotypes sourced from A) Siby, Mali and B) Kibos, Kenya when exposed to root exudate from RTx430 WT (*lgs1*-2), “wild type-like” null segregants (RTx430 L1,2 WT-L), insertion mutants of the *LGS1* gene (RTx430 L1,2 *LGS1*), a synthetic strigolactone analog (GR24), and water. L1,2 indicate mutant and null segregant pairs that originate from the same transformation event. Boxplots colored by *LGS1* allele of root exudate used as germination stimulants or controls. Asterisks denote statistically significant differences from the RTx430 WT by Student’s *t*-test. C) Source location (large white dots) of the two *S. hermonthica* ecotypes used in germination experiments. Denoted is each ecotype’s response to synthetic strigolactones, (+) 5-deoxystrigol (5DS) or (±) orobanchol, as reported in Bellis et al. 2020. \**\*P*<0.01, \*\*\*\**P*<0.0001 (*n*=15).

### Identifying potential transcription factor binding sites related to differential gene expression in lgs1-d plants

Using previously generated transcriptome data from Bellis et al. 2020, we identified impacted regulatory pathways in Macia *lgs1*-d plants (S_2_ generation). We used samtools v1.10 (Li et al., 2009) to extract the flanking genomic sequence of differentially expressed genes in roots and shoots separately. Specifically, we extracted the sequence 500 bp upstream of the gene’s start codon and the sequence 500 bp downstream of the gene’s stop codon, to capture differences in upstream and downstream regulatory coding regions. We then used STREME (Bailey, 2021) to determine significantly enriched motifs within the regulatory coding regions of differentially expressed genes compared to non-differentially expressed genes within each specific tissue (roots and shoots). We searched for overlap between enriched motifs and the JASPAR 2020 CORE (Anon) plant transcription factor binding sites database (Sandelin et al., 2004) with MEME suite’s online tool TOMTOM (Gupta et al., 2007).

### Growth conditions for mycorrhizal assay

We tested for differences in *LGS1* and Sb*CCD8b* functional and non-functional genotypes in the two genetic backgrounds (Maica, RTx430) under variable phosphate levels and grown with and without mycorrhizae. Plants were grown in a greenhouse in University Park, PA (16-hour photoperiod; day temperature minimum 21°C and maximum 28°C; night temperature minimum 21°C and maximum 26°C; relative humidity 20%) and subjected to three treatments: high phosphate without mycorrhizae (P+ AMF-), low phosphate without mycorrhizae (P-AMF-), and low phosphate with mycorrhizae (P-AMF+). The plants were grown in 10 inch “deepots” (Stuewe & Sons Inc., https://stuewe.com/product/2-5-x-10-heavyweight-deepot-cell/) in a mix of 90% sterile silica sand (Quickrete) and 10% kaolin clay (CB Minerals, https://cbminerals.com/kaolin/). After sowing, pots were watered with tap water every day until seven days after plant emergence. One-week post-emergence, plants were watered every other day with 200 mL water and treated with a supplemental fertigation of 1/3 Hoagland with different phosphate concentrations every other watering (every 4 days) until time of harvest. The phosphate level of the nutrient solution used at each fertigation was 333 μM for the high phosphate treatment (P+) or 33 μM for the low phosphate treatments (P-). Pots with a mycorrhizal treatment (AMF+) were inoculated with 50 mL of a sandy mix containing ∼20 spores/mL of *Rhizophagus intraradices* prior to planting. The spores used for inoculation were bulked in University Park, PA two months before the greenhouse experiment. Plants were harvested and destructive measurements were collected at two timepoints, 21 and 42 days after plant emergence (*n* = 7 per genotype, treatment, and time point; total *n* = 462).

The non-destructive measurements of light-saturated net carbon assimilation rate (steady state, measured via LI-6400), chlorophyll fluorescence and stomatal conductance (measured via LI-600 fluorometer/porometer), and shoot traits (plant height, vegetative phase change markers) were measured on greenhouse-grown plants prior to harvest dates. At the time of harvest (21 and 42 days) several destructive measurements were taken, and root tissue was collected for quantification of mycorrhizal colonization. Above ground tissue was collected and dried in an oven at 70°C for 72 hours for collection of dry biomass. As we could not measure root dry weight for plants grown in mycorrhizal treatments, fresh weight of the entire root system was measured. For the low phosphate non-mycorrhizal and mycorrhizal treatments, root systems were divided into biopsy cassettes (Fisher Scientific) for staining to quantify mycorrhizal colonization and confirmation of uncontaminated negative controls.

### Mycorrhizae counting assay

To test for differences in colonization of the fungal species *R. intraradices*, root samples of the mycorrhizal treatments stored in biopsy cassettes from both harvest time points (21 and 42 days) were stained (https://invam.ku.edu/staining-roots) and measured for relative mycorrhizal colonization (https://www2.dijon.inrae.fr/mychintec/Protocole/protoframe.html) (Phillips and Hayman, 1970; Koske and Gemma, 1989; McGONIGLE et al., 1990). At the time of harvest, cassettes were packed with ∼0.1 grams of root system and placed in 75% EtoH until time of staining. Roots were cleared using 10% KOH that was brought to boiling and then removed from the heat prior to adding cassettes. Cassettes containing root samples were left in the hot KOH solution for 10 minutes. Samples were then washed five times using diH_2_O and acidified by being immersed in a 2% HCl solution for 20 minutes at room temperature. After acidification, roots were stained from incubation in a mixture of 1:1:1 water, glycerin, and lactic acid containing 0.05% direct blue at room temperature for at least 12 hours. Stained samples were again rinsed with diH_2_O five times and stored suspended in diH_2_O in a fridge at 15°C for one week to allow for excess leaching of the stain prior to quantification of mycorrhizae.

Relative mycorrhizal colonization was quantified using a modified version of the grid line intersect method (Newman, 1966; Tennant, 1975; Giovannetti and Mosse, 1980). Stained roots were randomly dispersed in a nine-cm diameter petri plate with an acetate sheet containing 10×10 grid lines under the petri plate. Using an Amscope SM-2, intersections where a root crossed a grid line were designated as either non-mycorrhizal or colonized. The presence of any fungal structure (intracellular hyphae, vesicle, arbuscule, spore) resulted in the intersection being scored as “colonized”. To assess the relative abundance of mycorrhizae in each sample, 100 intersections were scored, and relative colonization was calculated as a percentage.

### Collection of root phenotypes

Parameters of root system architecture and anatomy were collected for plants grown in the high phosphate no mycorrhizae treatment at the 42 days after plant emergence time point. To characterize architecture, entire root systems were scanned on an Epson scanner and processed in RhizoVision Explorer v2.0.3 (Seethepalli and York, 2020; Seethepalli et al., 2021) using algorithms described by (Seethepalli et al., 2021). For root anatomy, we collected two representative axial roots from node three and excised two four-cm root samples five to nine cm from the most basal portion of the whole root sample. Root samples were stored in 75% ethanol until being sectioned by laser ablation tomography (LAT; (Strock et al., 2019)). In LAT, a sample is moved via an automatic stage towards a 355-nm Avia 7000 pulsed laser and ablated in the focal plane of a camera. A Canon T3i camera with a 53micro lens (MP-E 65 mm) was used to capture images of the root cross-section. Two representative images for each root sample sectioned one to three cm apart and were saved for later image analysis with Rootscan (https://plantscience.psu.edu/research/labs/roots/methods/computer/rootscan) (Burton et al., 2012). For each plant, anatomical phenotypes were averaged across images collected for each nodal root and reported for the two sampled roots.

### Photosynthesis, stomatal conductance, and density of stomata

All steady state photosynthesis measurements were measured from plants grown without nutrient limitation using a Li 6400 portable photosynthesis machine (Li Cor Environmental). Prior to collecting gas exchange measurements, leaves were acclimated to the starting chamber conditions and the first data point was logged after CO_2_ and transpiration rates reached steady state. For all measurements, a constant leaf temperature of 25°C, light level of 1800 µmol m^−2^ s^−1^, and a flow rate of 400 µmol s^−1^ were maintained. Light saturated photosynthetic rate (*A*_sat_) was measured at a reference CO_2_ level of 400 ppm. *A*_sat_ was recorded every 20 seconds for two minutes and reported as the average across all data points logged.

The Li-600 Porometer/Fluorometer machine (Li Cor Environmental) was also used to measure stomatal conductance and parameters of dark (leaves were kept in the dark for a minimum of 30 minutes) and light-adapted chlorophyll fluorescence. For all samples, light and dark-adapted measurements were collected on the same leaves with the fixed flow rate set to high (150 µmol s^−1^) and a flash intensity of 6000 μmol m^-2^ s^-1^.

We further collected CO_2_ response curves, steady state photosynthesis, and ran untargeted metabolomics for the previous S_2_ generation of Macia *lgs1*-d CRISPR deletion mutants using plant material as described in (Bellis et al., 2020) in a separate greenhouse experiment. Here, we also collected steady state photosynthesis for RTx430 *ccd8b.* Plants were grown in a media mix that consisted of 80% commercial sand (Quickrete) and 20% turface (LESCO turface, https://www.siteone.com/en/088782-lesco-turface-all-sport-soil-conditioner-infields-50-lb/p/23795) for 16-hour days. After germination, plants were watered with 100 mL of water every other day and supplementally fertigated with 100 mL half strength Hoagland once a week starting at 7 days after plant emergence. CO_2_ response curves were built by measuring net photosynthetic rate (*A*_net_) at the reference CO_2_ levels of 50, 100, 200, 300, 400, 600, 800, 1000 ppm. Across all CO_2_ levels, a constant leaf temperature of 25°C, light level of 1800 µmol m^−2^ s^−1^, and a flow rate of 400 µmol s^−1^ were maintained. Leaves were given a chance to acclimate to the chamber CO_2_ conditions for at least two minutes before data logging at each CO_2_ level. We used methods described in (Zhou et al., 2019) to fit *A*_net_ *C_i_* curves and to determine apparent *V*_cmax_ and apparent *J_max_* using procedures of (Sharkey et al., 2007), accounting for carbonic anhydrase activity.

Leaves used in gas exchange measurements were also analysed to determine stomatal density. Here, we collected three non overlapping fields of view from comparable regions on the abaxial and adaxial surface. Images were captured using an AmScope MU1400 digital camera mounted on a compound microscope equipped with a 20×/0.25 finite objective (F-LD 20/0.25 160/2) and the camera was connected through a 0.5× C mount adapter, resulting in an effective magnification of approximately 10×. Number and area of stomata were quantified using ImageJ (Schneider et al., 2012). All area measurements were converted from pixel units to micrometers using the calibrated pixel size (≈ 0.14 µm px^-1^).

### Metabolomics

To quantify differences in abundance of small molecules for sorghum mutants of Sb*CCD8b* and *LGS1*, root and shoot tissue samples were taken for *n* = 5 plants of the genotypes RTx430 WT, RTx430 *ccd8b*, Macia WT, and the S_2_ generation of Macia *lgs1*-d deletion lines (Bellis et al. 2020) and for evaluation with untargeted metabolomics. Root and shoot samples were immediately flash-frozen separately in liquid nitrogen and transferred to a 1.5 mL Eppendorf tube for further processing. Samples were then lyophilized at -80°C overnight before being ground to a fine powder by placing a metal ball in each Eppendorf tube and shaking in a TissueLyser II (Qiagen) for 5 minutes at 25 Hz.

All solvents were chromatography grade and acquired from VWR (Radnor, PA, U.S.A.) or Sigma Aldrich (St. Louis, MO, U.S.A.). Metabolites were extracted using 1 mL 80% MeOH + 0.1% formic acid for every 100 mg of powdered sample. Samples were vortexed for 10 seconds and sonicated for five minutes. Then, each sample was centrifuged at 10,000 rpm for 10 minutes and the supernatant was transferred to a 1 mL glass scintillation vial. The extraction process was repeated with the same tissue and the supernatants were combined before being air-dried to completion in a fume hood. All samples were reconstituted at a concentration of 1 mg/mL in LC-MS grade methanol with 1 uM chlorpropamide as an internal standard. A Vanquish Duo UHPLC system connected to a Thermo Fusion Lumos Orbitrap Mass Spectrometer (Thermo Fisher Scientific, Waltham, MA) was used for metabolomics analysis with a Waters Acquity UPLC BEH C18 (21.x150 mm x 1.7 um) column. The flow rate was 0.1 mL/min at 55°C. Solvent A was 0.1% formic acid (v/v) in LC-MS water and solvent B was 0.1% formic acid (v/v) in LC-MS acetonitrile. The mobile phase gradient of solvent B was: 3% for 0.01 minutes, 45% for 10 minutes, 75% for 2 minutes, 100% for 4.5 minutes, and 3% for 0.2 minutes. A 2 ul injection was used for all samples.

Mass spectrometry was conducted using an electrospray ionization source with a positive ion spray voltage of 3500 V, sheath gas pressure of 25 Arb, auxiliary gas pressure of 5 Arb, ion transfer temperature of 275°C, and vaporizer temperature of 75°C. MS1 data was acquired with an Orbitrap resolution of 120,000, scan range of 100-1000 Da, and RF lens of 50% in the profile mode. MS2 data was collected in a data-dependent manner using intensity (2.5E4) and dynamic exclusion (ions excluded after 1 detection for 30 seconds).

All .raw mass spectral files were converted to .mzML using MSconvert (Proteowizard) and then loaded into MzMine3.1 (Schmid et al., 2023). Root and shoot files were analyzed separately. Supplemental Table 5 outlines the MZMine3 processing parameters. Following processing in MzMine3.1, all features with a peak area less than 5-fold the intensity of the metabolite concentration in the blanks were removed. The peak area was Hellinger transformed and auto-scaled. To make volcano plots, the compounds’ *P*-values and log_2_fold changes were calculated between each mutant and its corresponding wild type. Compounds with a log_2_fold change greater than 0.6 and a *P*-value less than 0.05 were labeled as “up-regulated” and those with a log_2_fold change less than 0.6 and a *P*-value less than 0.05 were labeled as “down-regulated.” Key compounds were annotated by comparing their MS2 profiles to potential matches found via molecular networking performed using the Global Natural Products Social Molecular Networking (GNPS) library search feature (Wang et al., 2016). To compare statistical differences between mutants and their respective wild type lines, a Permutational Multivariate Analysis of Variance (PerMANOVA) was performed using the R/vegan (Oksanen et al., 2022).

### Quantification of chlorophyll content and light harvesting proteins

We assessed if chlorophyll content (Chl*a*/Chl*b*), and concentration of photosystem proteins (ratio of PSAA/CP43 and LHCB1/CP43) were altered in sorghum genotypes segregating for functional Sb*CCD8b* and *LGS1*. Plants used in this quantification were watered every day for the first week and then every other day until time of harvest. At every other watering, plants were supplementally fertigated with 1/3 Hoagland solution. At the time of harvest, five-week-old plants were flash frozen in liquid nitrogen. Chlorophylls were extracted from leaf tissue and thylakoid samples in acetone 80% and the absorption spectra were analyzed using the Cary and fitted as previously described (Croce et al., 2002).

Total proteins were extracted by grinding the frozen leaf tissue in a mortar with the BufferA/4 from (Schägger, 2006) (3% SDS (wt/vol), 1.5% mercaptoethanol (vol/vol), 7.5% glycerol (wt/vol), 0.012% CoomassieblueG-250 (Serva), 37.5mMTris/HCl (pH7.0)) supplemented with protease inhibitor (cOmplete™ Protease Inhibitor Cocktail (Roche)). Samples were then centrifugated and the supernatant was stored at -20°C. Immunoblotting was performed as described in (Schägger, 2006). All primary antibodies, PsaA(AS06172), PsbC(AS111787), and LHCB1(AS01004), were purchased from Agrisera, Sweden. Chemilluminescence was collected by a LAS 4000 Image Analyzer and analyzed with the software ImageJ.

### Statistical analysis

For statistical analysis of the impact of exudate on *Striga* germination, we built negative binomial generalized linear models using R/MASS::glm.nb (Venables and Ripley, 2002). Nested models accounted for transformation event and *S. hermonthica* ecotype as effects, with one model additionally including an allele term. The two models were compared using a likelihood ratio test and significance of the *LGS1* allele effect was assessed using a chi-square distribution.

For all other collected phenotypes, we tested for the significance of *LGS1* deletion (*lgs1*-d) and *LGS1* insertion alleles by fitting linear mixed effects models using R/lme4::lmer (Doran et al., 2007). Full models included the fixed effect of allele and the random effect of transformation event and a reduced model excluded the fixed effect allele term. To determine if the inclusion of the allele term improved model fit, the full and reduced models were assessed with a likelihood ratio test implemented with the R ‘anova’ function and significance was determined based on the resulting chi-square statistic and associated *P*-value. Separate models were built for plants of the Macia and RTx430 background. For Sb*CCD8b* deletions, we used a student’s *t*-test to assess significance of the deletion allele. R statistical software (R Core Team, 2022) was used throughout for data processing and statistical analysis.

## Results

### Presence-absence variation of LGS1 and Sb3500 in natural populations and predicted strigolactone profiles

Previous studies demonstrate substantial structural variation across the *LGS1* genomic region in sorghum (Gobena et al., 2017; Bellis et al., 2020; Maina et al., 2025). To contextualize our experimental genotypes, we assessed presence-absence variation of two genes that impact strigolactone profiles, *Sb3500* (*Sobic.005213500*) and *LGS1* (*Sobic.005213600*), in both our experimental lines and natural populations, and used published functional data to infer expected strigolactone profile (Table 2).

Genotypes with putatively functional copies of both genes (e.g. Macia WT) were the most common in sorghum landraces (*N* = 668) and are expected to produce predominantly 5DS type strigolactone (Gobena et al., 2017; Yoda et al., 2023). In contrast, genotypes with *LGS1* deletion alleles and presence of *Sb3500* (e.g. Macia *lgs1*-d, *lgs1*-3) are expected to primarily exude orobanchol (Gobena et al., 2017; Bellis et al., 2020) and occur in sorghum landraces at low frequency (*N* = 15). The RTx430 WT carries the natural *lgs1*-2 deletion allele which spans both *Sb3500* and *LGS1*. This absence at both these genes also occurs at a low frequency in sorghum landraces (*N* = 20), and orobanchol is the expected predominant strigolactone in exudate (Gobena et al., 2017; Yoda et al., 2021). Importantly, the RTx430 *LGS1* insertion mutant represents a distinct and extremely rare haplotype (*N* = 1) in which *LGS1* is present in the absence of *Sb3500.* Transient expression of *LGS1* in *E. coli* incubated with a strigolactone precursor produced 5DS and 4DO in approximately equal amounts, while co-expression with *Sb3500* resulted in the stereoselective biosynthesis of 5DS in *Nicotiana benthamiana* (Yoda et al., 2023). As such, we hypothesize that the RTx430 *LGS1* insertion genotype, absent of functional *Sb3500*, produces an intermediate strigolactone profile of 5DS and 4DO and is distinct from the strigolactone profile of RTx430 WT (orobanchol), and the vast majority of genotypes with the *LGS1* gene present (high 5DS) (Mohemed et al., 2018).

### Inserting functional LGS1 into a naturally loss-of-function background restores Striga hermonthica germination

We sought to confirm that the new mutants with functional *LGS1* inserted into the RTx430 background changed strigolactone profiles in a manner consistent with the deletion effects (Q1). We tested the germinability of two *S. hermonthica* ecotypes known to be differentially sensitive to (+) 5DS and (±) orobanchol (Bellis et al., 2020)(Figure 1C). Root exudate collected from the RTx430 *LGS1* insertion mutants induced significantly more germination of both *S. hermonthica* ecotypes as compared to RTx430 WT and WT-L null segregants with the *lgs1*-2 natural deletion allele (χ^2^ *=* 53.95, *P <* 0.001, likelihood ratio test for fixed effect of *LGS1* deletion; Figure 1A-B). The magnitude of RTx430 *LGS1* effect on *S. hermonthica* germinability differed between the two ecotypes, supporting that there is genetic variation in parasite sensitivity to host-derived strigolactone profiles of root exudates. For the *S. hermonthica* ecotype from Siby, Mali, average germination was <0.02% when treated with exudate from RTx430 WT and WT-L (*lgs1*-2) plants and 6.5% when treated with exudate from the RTx430 *LGS1* insertion mutant plants (Figure 1A). The *S. hermonthica* ecotype from Kibos, Kenya had slightly higher germinability, and an average germination rate of <1% when treated with RTx430 WT and WT-L (*lgs1*-2) exudate and 10.56% when treated with exudate from RTx430 *LGS1* (Figure 1B). Variation in parasite sensitivity to strigolactone profiles for the same two ecotypes was also reported by Bellis et al. (2020), in which Macia *lgs1*-d mutants had lower germinability of the parasite compared to Macia WT, with this trend being weaker for the Kenyan ecotype. Notably, this Kenyan *S. hermonthica* ecotype has a similar sensitivity to (+) 5DS and (±) orobanchol (Figure 1C) (Bellis et al., 2020). Here, we observe little to no germination for both *S. hermonthica* ecotypes when exposed to exudates from RTx430 WT and WT-L plants, suggesting that the *lgs1*-2 natural deletion allele may not only shift strigolactone profiles in matter that does not strongly induce germination (i.e. orobanchol), but also reduce the overall strigolactone in root exudate, as exemplified by low germination in the Kenyan population. Together, insertion of functional *LGS1* into a natural loss-of-function background, RTx430 (*lgs1*-2), changes strigolactone profiles in a manner that increased the germination of *S. hermonthica* likely due to an increase of 5DS and a larger quantity of strigolactone in root exudate.

### Pathways involved in gene regulatory effects of LGS1 deletion

Bellis et al. (2020) reported that deleting functional *LGS1* from the otherwise functional Macia background resulted in substantial changes in gene expression. In *LGS1* knockout plants (Macia *lgs1*-d), 505 differentially expressed genes (DEGs) were identified in roots (244 down-regulated, 261 up-regulated) and 2,167 DEGs were identified in shoots (917 down-regulated, 1,250 up-regulated). To identify regulatory pathways that relate to differences in gene expression (Q2), we searched for sequence motifs in putative promoter regions of *lgs1*-d DEGs using STREME (Bailey, 2021). Within putative promoter regions of Macia *lgs1*-d root DEGs, we discovered three enriched sequences that overlapped with 16 known motifs, and within shoot DEGs we found six enriched sequences that overlapped with 92 known motifs. We next sought to match enriched motifs identified in putative promoter regions of DEGs to known transcription factor binding site (TFBS) motifs. In roots, the most commonly discovered motif (*GACCGTCCATC*) was not significantly enriched in DEGs (*P* = 0.48), indicating that it is broadly prevalent across expressed root genes, rather than specific to differential expression. This motif overlapped with members of six known TFBS families (Table S1), including ethylene-responsive element-binding factors (ERFs) and Squamosa Promoter Binding Protein-Like (SPL) genes (overlap of discovered motif with TFBS sequences, *P* < 0.004), which are involved in abiotic stress response and lateral root development and nodulation, respectively (Fujimoto et al., 2000; Yu et al., 2015; Nasrollahi et al., 2022). Together, the identified root TFBS motifs reflect regulatory patterns among expressed genes in roots; however, these motifs were not significantly associated with the regulatory variation underlying differential expression in Maica *lgs1*-d roots. For shoots, the most common discovered motif (*CCGGCAGCG*) was significantly enriched in 41 DEGs (*P* = 0.002, Table S2). Of known TFBS significantly overlapping motifs enriched in promoters of shoot DEGs, the majority belonged to the APETALA 2/ethylene-responsive element binding proteins (AP2/EREBP) family (overlap of discovered motif with TFBS sequences, *P* < 0.02), which are key regulators of leaf and floral development and participate in response to biotic and environmental stress (Riechmann and Meyerowitz, 1998). AP2/ERFs regulate numerous target genes, suggesting that deleting *LGS1* in a normally functional background may lead to broad pleiotropic effects, and stress and growth responses may be particularly disrupted in Macia *lgs1*-d.

### Reduced CO_2_ assimilation rate in strigolactone biosynthesis deletion mutants

Strigolactones have been demonstrated to be positive regulators of light-harvesting genes in tomato (Mayzlish-Gati et al., 2010) and Bellis et al., (2020) reported significant downregulation of photosystem I, photosystem II, and light-harvesting genes in Macia *lgs1*-d knockout lines. To assess whether this transcriptional signal translates into downstream phenotypic effects, we evaluated mutant plants for differences in assimilation rate, chlorophyll content, stomatal conductance, stomatal density, and the relative abundance of core light-harvesting proteins (Q3).

In the strigolactone-deficient RTx430 *ccd8b* line, light-saturated net CO_2_ assimilation rate (*A*_sat_) was significantly reduced compared to RTx430 WT (t(10) = 2.36, *P* = 0.040, difference of genotypes by Student’s *t*-test; Figure 2D). This reduction may reflect changes in stomatal regulation and/or photosynthetic capacity. RTx430 *ccd8b* mutants had reduced stomatal density on both abaxial and adaxial leaf surfaces (t(34) = -2.1684, *P* = 0.037, difference of genotypes by Student’s *t*-test; Figure S2A-B). Under non-limiting nutrient conditions, stomatal conductance did not significantly differ between RTx430 *ccd8b* and WT plants, although it was elevated in the mutant earlier in development (Figure S2C-D). RTx430 *ccd8b* also had lower values for chlorophyll a/b ratio (Chl*a*/Chl*b*), the ratio of antenna complexes to PSII core proteins (LHCB1/CP43), and the ratio of PSI to PSII core proteins (PSAA/CP43) potentiating physical differences in photosynthetic machinery relative to the RTx430 WT; however, these differences were not statistically significant (Figure S3). Taken together, these results suggest the contribution of stomatal traits for reduced carbon assimilation in RTx430 *ccd8b*, although the roles of stomatal versus biochemical limitations remain unresolved.

**Figure 2.**
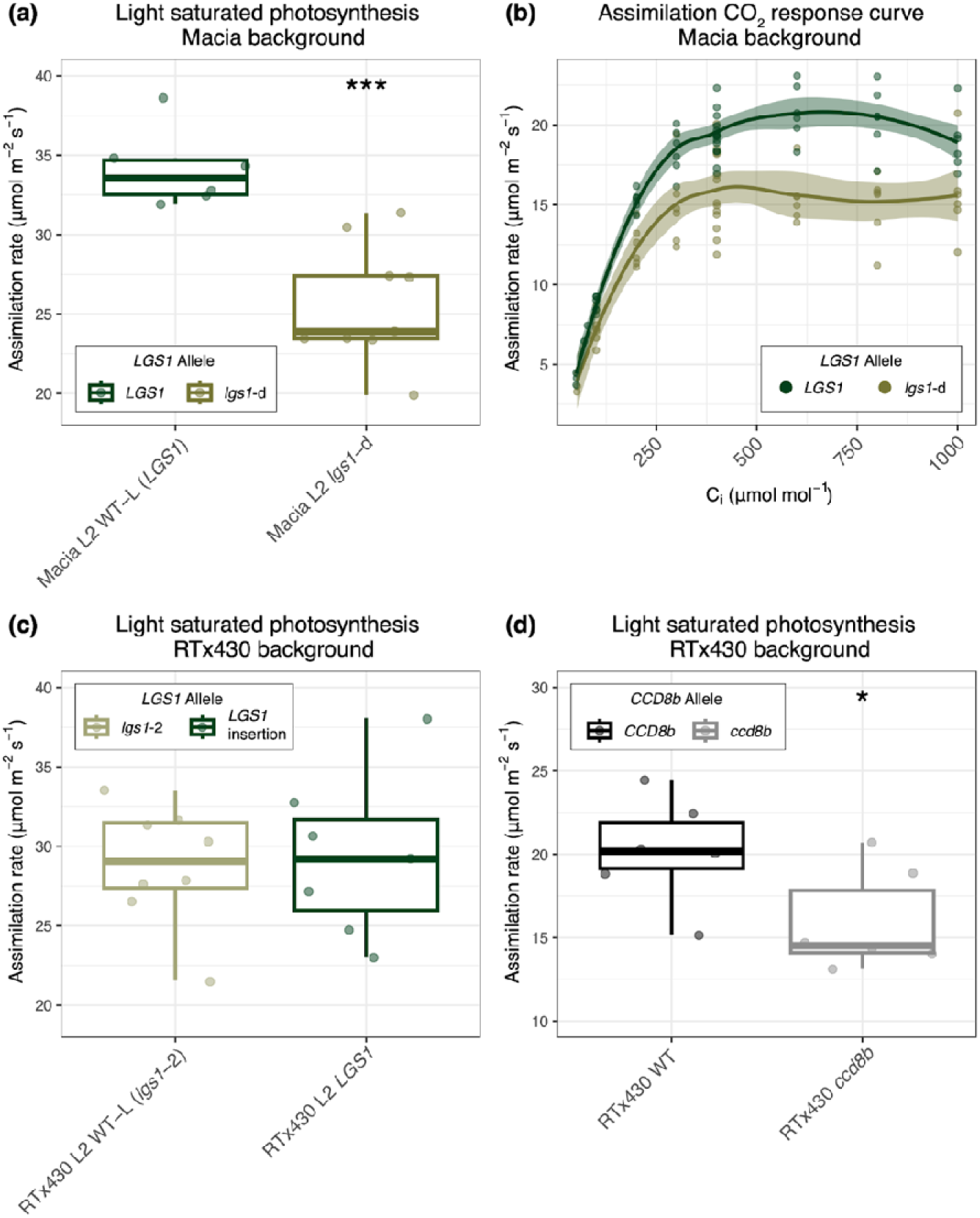
Light saturated assimilation rate is reduced when strigolactone biosynthesis genes are knocked out of normally functional backgrounds. A) Net carbon assimilation rate of *LGS1* deletion (*lgs1*-d) mutants and null segregants (WT-L) of the same transformation event (L2) generated in the Macia background. B) Assimilation rate of the Macia WT and *LGS1* deletion (*lgs1*-d) mutants at varying levels of *C_i_*. Data is collected with mutants of the S_2_ generation, see Figure S4 for net carbon assimilation rate of S_2_ generation Macia *lgs1*-d. C) Net carbon assimilation rate of mutants with functional *LGS1* inserted and “wild type-like” null segregants (WT-L *lgs1*-2) of the same transformation event (L2) generated in the RTx430 background. D) Net carbon assimilation rate of *ccd8b* mutants and WT in the RTx430 background. Asterisks denote a significant *P*-value comparing each mutant and wild type (WT or WT-L) comparison by Student’s *t*-test. \**P*<0.05, \*\*\**P*<0.001 (*n*=6).

For mutants of the *LGS1* gene, the rate of light-saturated photosynthesis (*A*_sat_) was on average 25% lower in the deletion mutant (Macia *lgs1*-d; ^2^(1) = 6.8157, *P* = 0.009, likelihood ratio test for fixed effect of *LGS1* deletion; Figure 2A, Figure S4A), but was not significantly different in the insertion mutant (RTx430 *LGS1*; Figure 2C) relative to each WT comparison. Despite a reduced stomatal density on both abaxial and adaxial leaf surfaces for Macia *lgs1*-d (Figure S4D-E), stomatal conductance was not significantly different from the WT (Figure S4F), suggesting differences in Macia *lgs1*-d *A*_sat_ is not attributable to stomatal constraints. We next assessed if reductions in *A*_sat_ for Macia *lgs1*-d reflected changes in photosynthetic biochemistry, and additionally examined the maximum Rubisco carboxylation rate (*V*_cmax_) and the maximum electron transport rate for ribulose-1,5-bisphosphate regeneration (*J*_max_), estimated from *A*_net_–*C_i_* curves, along with fluorescence-derived traits (Fv/Fm, maximum quantum efficiency of photosystem II (PSII); ΦPSII, quantum yield of PSII; non-photochemical quenching (NPQ); electron transport rate (ETR)), and the relative abundance of core light-harvesting proteins.

In Macia *lgs1*-d lines, *V*_cmax_ was significantly reduced ( ^2^(1) = 5.00, *P* = 0.025, likelihood ratio test for fixed effect of *LGS1* deletion; Figure S4B) and apparent *J*_max_ showed a similar but non-significant trend toward reduction ( ^2^(1) = 3.25, *P* = 0.07, likelihood ratio test for fixed effect of *LGS1* deletion; Figure S4C). In contrast, fluorescence traits did not differ significantly between genotypes. Reductions in *V*_cmax_ and *J*_max_ are consistent with altered photosynthetic capacity, and may relate to differences in nitrogen allocation or photosystem investment (Walker et al., 2014). Despite reduced assimilation rates of Macia *lgs1*-d, the overall quantification of chlorophyll content and relative abundance of antenna and core photosystem proteins revealed no detectable differences between *lgs1*-d mutants and WT or WT-L lines in the Macia background (Figure S3). Together, the reductions in net carbon assimilation rate may be attributable to photosynthetic efficiency, such as Rubisco activity or activation state (as indicated by reduced *V*_cmax_ in Macia *lgs1*-d), rather than structural changes to the photosynthetic machinery. Notably, differences in assimilation rate between Macia and RTx430 functional and non-functional *LGS1* suggest *LGS1* effects as dependent on genetic background (Q4), and underscores the importance of evaluating QTL in multiple genetic backgrounds, when possible.

### Mutation of strigolactone genes impacts the metabolome

Previous research has reported differences in the metabolic profiles of unrelated, non-isogenic sorghum genotypes that segregate for *LGS1* (Kawa et al., 2021) and some metabolites, especially flavonols, are observed to have strigolactone-dependent accumulation (*A. thaliana* (Walton et al., 2016)). We sought to identify any differences in the root and shoot metabolome of Sb*CCD8b* and *LGS1* knockout lines through untargeted metabolomics using ultra-high performance liquid chromatography-MS/MS (UHPLC-MS/MS). For both deletion mutants, we observed differences in metabolite intensity compared to wild type (Figure S5). Most of these shifts were in low-abundance compounds that we were unable to identify via library matching and molecular networking (Table S3, S4) and did not dramatically change the overall profile of the metabolome (PerMANOVA of root and shoot untargeted metabolomic profiles, both genes, *P* > 0.5; Figure S6). An exception was a major shift in the production of a disaccharide that we have tentatively identified as maltose, which was increased in the roots and shoots of RTx430 *ccd8b* and had variable shifts in the roots and shoots of Macia *lgs1*-d (Figure S5).

For RTx430 *ccd8b*, the overall base peak chromatogram (BPC) intensity of both roots and shoots was higher than the RTx430 WT, indicating elevated total metabolite production when Sb*CCD8b* is deleted (Figure S5A-B). The majority of RTx430 *ccd8b* differentially accumulated metabolites were up-regulated in the roots (Figure S7A). Of the RTx430 *ccd8b* up-regulated root metabolites, we putatively identified dihydroorotate which has been previously reported as up-regulated in drought-sensitive sorghum (Li et al., 2022) and is consistent with observations of strigolactone biosynthesis mutants’ increased sensitivity to drought (Van Ha et al., 2014). Further, dihydroorotate dehydrogenase (*DHODH*) is an enzyme involved in pyrimidine biosynthesis and its overexpression in rice both increased *DHODH* activity and enhanced plant tolerance to salt and drought stresses (Os*DHODH1*; (Liu et al., 2009)).

Though we did not detect significant differences between the overall metabolome of Macia *lgs1*-d as compared to Macia WT Figure S6A-B, differentially accumulated metabolites were more down-regulation in the roots (Figure S7C,E,G) and up-regulated in the shoots (Figure S7D,F,H). A metabolite with a putative annotation of 2,6-Dimethylpyrazine, a natural product belonging to the class of pyrazine-class volatile organic compounds (VOCs), was significantly down-regulated in roots of Macia *lgs1*-d (Table S4). Though direct studies on 2,6-dimethylpyrazine are limited, it is assigned to a pyrazine-class of VOCs that can activate defense pathways (Patel et al., 2021), and act as signal molecules perceived by plants (Khashaveh et al., 2025). Its down-regulation in the roots of Macia *lgs1*-d once again suggests alteration to stress response when *LGS1* is knocked out of a normally functional background. For both Sb*CCD8b* and *LGS1* loss-of-function mutants, differences in low-abundance metabolites suggesting some metabolic pathways were affected, though the most abundant metabolites remained largely unchanged.

### Strigolactone synthesis mutations impact root architecture and anatomy

Strigolactones are well established to play critical roles in modulating root architecture and more recently have been linked to root anatomical differences to optimize vessel element formation (Zhao et al., 2025) and in aerenchyma formation (Shrestha et al., 2026). Consistent with previous reports by Hoa et al., (2023), which demonstrated RTx430 *ccd8b* to have fewer root tips and overall smaller root systems, we also report RTx430 *ccd8b* plants had significantly smaller root systems (t(14) = 4.54, *P* < 0.001, difference of genotypes by Student’s t-test; Figure 3C). Additionally, we found that RTx430 *ccd8b* root systems had significantly more nodal roots (t(14) = -3.17, *P* = 0.007, difference of genotypes by Student’s t-test; Figure 3D), with roots that were smaller in cross section area (t(23) = -3.43, *P* = 0.002, difference of genotypes by Student’s t-test; Figure 3E), and had altered root anatomy (Figure 3A-B). Most notably, RTx430 *ccd8b* aerenchyma area (air space in the root cortex) was significantly reduced (t(17) = -2.60, *P* = 0.019, difference of genotypes by Student’s t-test; Figure 3H) and despite maintaining the same number of metaxylem vessels, trended towards reduction in total metaxylem vessel area (t(9) = - 1.71, *P* = 0.123, difference of genotypes by Student’s t-test; Figure 3F-G).

**Figure 3.**
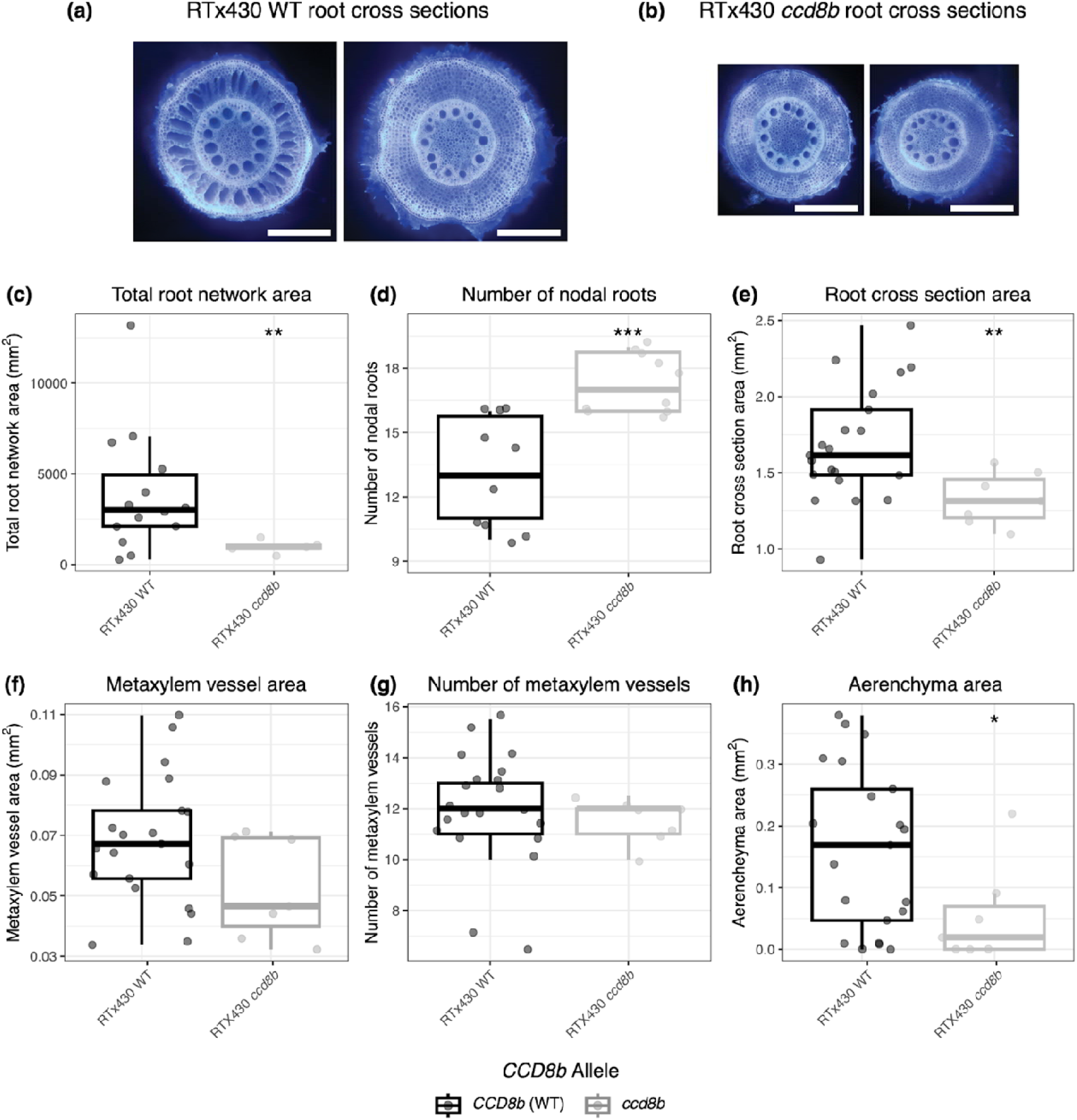
Root Architecture and anatomy is altered in mutants with Sb*CCD8b* deleted. Two representative root cross sections captured by laser ablation tomography (LAT) for the RTx430 background A) WT and B) *ccd8b* mutants. The scale bar in each photo is 0.5 mm. Boxplots compare the RTx430 WT (black) and *ccd8b* mutants (gray) for the root architecture phenotypes of C) total root system network area D) number of nodal roots and the root anatomical traits of E) root cross-section area F) total metaxylem vessel area G) number of metaxylem vessels H) total aerenchyma area. Asterisks denote a significant *P*-value comparing WT and mutant by Student’s *t*-test. \**P*<0.05, \*\**P*<0.01, \*\*\**P*<0.001.

Macia *lgs1*-d maintained a greater number of metaxylem vessel elements ( ^2^(1) = 5.67, *P* = 0.017, likelihood ratio test for fixed effect of *LGS1* deletion; Figure S8E) with individual elements that were smaller in size ( ^2^(1) = 4.20, *P* = 0.04, likelihood ratio test for fixed effect of *LGS1* deletion; Figure S8F). For the aerenchyma area trait, the Macia *lgs1*-d mutant had a reduced formation of airspace in the cortex as compared to the Macia WT and WT-L null segregants, which showed greater plasticity in the formation of nodal root aerenchyma ( ^2^(1) =1.26, *P* = 0.26, likelihood ratio test for fixed effect of *LGS1* deletion; Figure S8G). Differences in root architecture anatomy for the RTx430 mutants with functional *LGS1* inserted were not as pronounced and not consistent with observations in Macia *lgs1*-d mutants. RTx430 *LGS1* mutant had a greater number of nodal roots as compared to the RTx430 WT (t(43)=-2.715*, P* = 0.001, likelihood ratio test for fixed effect of *LGS1* deletion; Figure S9B). All other root traits were not significantly different. Interestingly, both genotypes with functional strigolactone genes knocked out (RTx430 *ccd8b*, Macia *lgs1*-d) had reduced aerenchyma area and is consistent with recent reports by (Shrestha et al., 2026) that strigolactones play a role in aerenchyma formation.

### Strigolactone synthesis mutants grown in nutrient and mycorrhizal treatments

We hypothesized the alteration of sorghum strigolactone profiles from loss-of-function at *LGS1* that reduce parasite stimulation (Figure 1, Bellis et al. 2020), would also be coupled with trade-offs in plant growth and interactions with arbuscular mycorrhizal fungi (AMF). As strigolactones mediate rhizosphere signaling for symbiosis, we predicted that loss-of-function at Sb*CCD8b* and *LGS1* would impair mycorrhizal establishment (Q1). To test this, all genotypes (Table 1) were evaluated under three conditions: low phosphate without mycorrhizae, low phosphate with mycorrhizae, and high phosphate without mycorrhizae (Figure 4).

**Figure 4.**
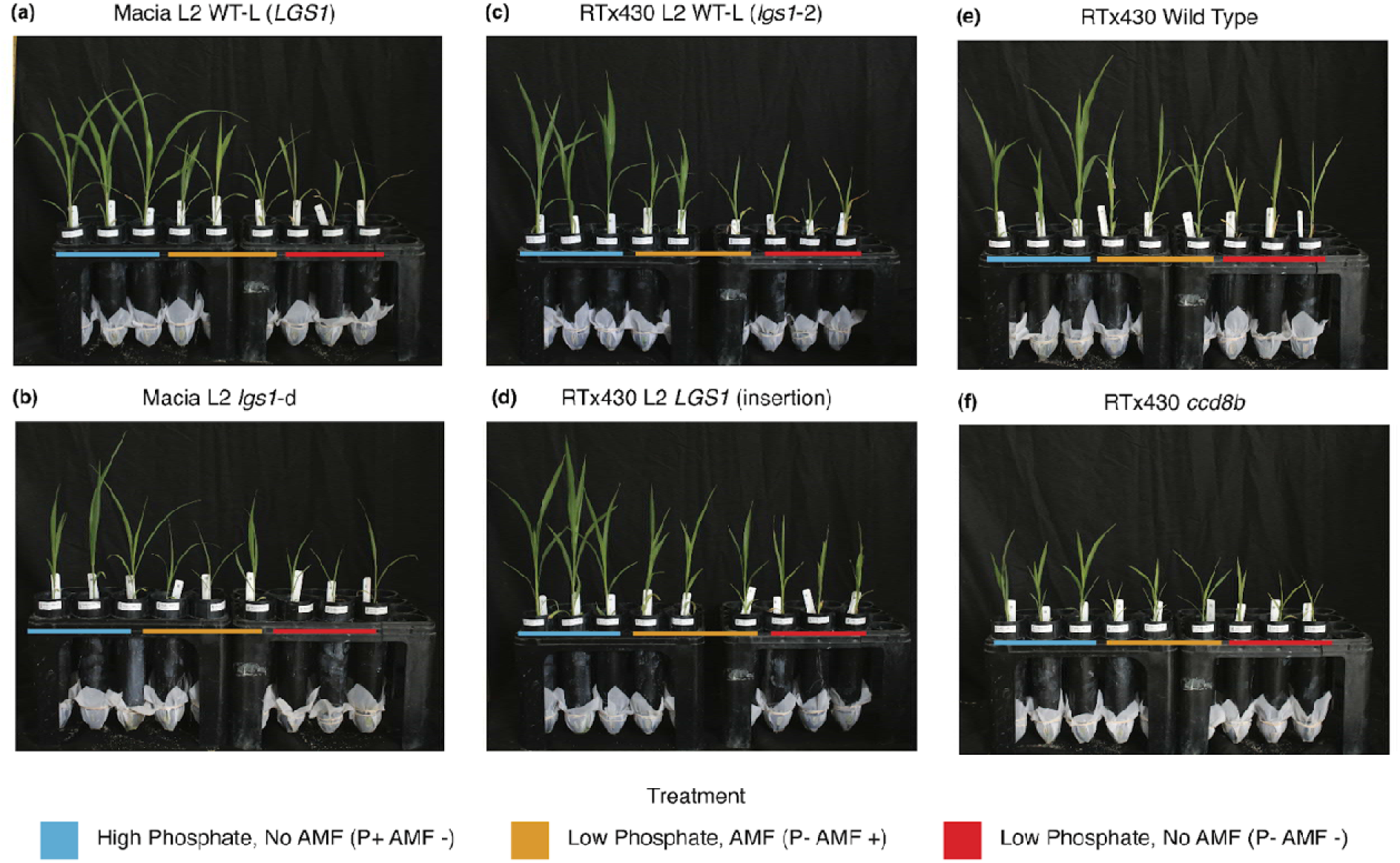
Representative images of greenhouse grown plants taken at 28 days after plant emergence. For a genotype (denoted above image), three replicates grown in the three treatments are included in a photo. The color bar corresponds to the treatment a plant was grown in. Treatments are high phosphate and no arbuscular mycorrhizal fungi (P+ AMF-; blue), low phosphate and arbuscular mycorrhizal fungi (P-AMF+ ; orange), and low phosphate and no arbuscular mycorrhizal fungi (P-AMF-; red). A,B) Comparison of “wild type -like” null segregants (WT-L) and deletion mutants of the *LGS1* gene (*lgs1*-d) in the Macia background. C,D) Comparison of “wild type -like” null segregants (WT-L *lgs1*-2) and insertion mutants of the *LGS1* gene RTx430 background. E,F) Comparison of WT and deletion mutants of the Sb*CCD8b* gene (*ccd8b*) in the RTx430 background. Description of genotypes are reported in Table 1.

We observed strong treatment effects, and plants were the tallest and largest in the high phosphate treatment (P+ AMF-, Figure 5A). The application of phosphate (P+) and mycorrhizal inoculation (AMF+) each independently increased plant height, shoot weight, root weight, a SPAD-based measurement of chlorophyll concentration, and stomatal conductance, as compared to the low phosphate non-mycorrhizal treatment (P-AMF-)(Figure 5A-E). For all genotypes, chlorophyll concentration and stomatal conductance were the lowest in the low phosphate no mycorrhizae treatment (P-AMF-), as compared to the treatments with high phosphate (P+, AMF-) and with mycorrhizae (P-AMF+) (Figure 5D-E), indicating that the low phosphate no mycorrhizae treatment (P-AMF-) was indeed stressful.

**Figure 5.**
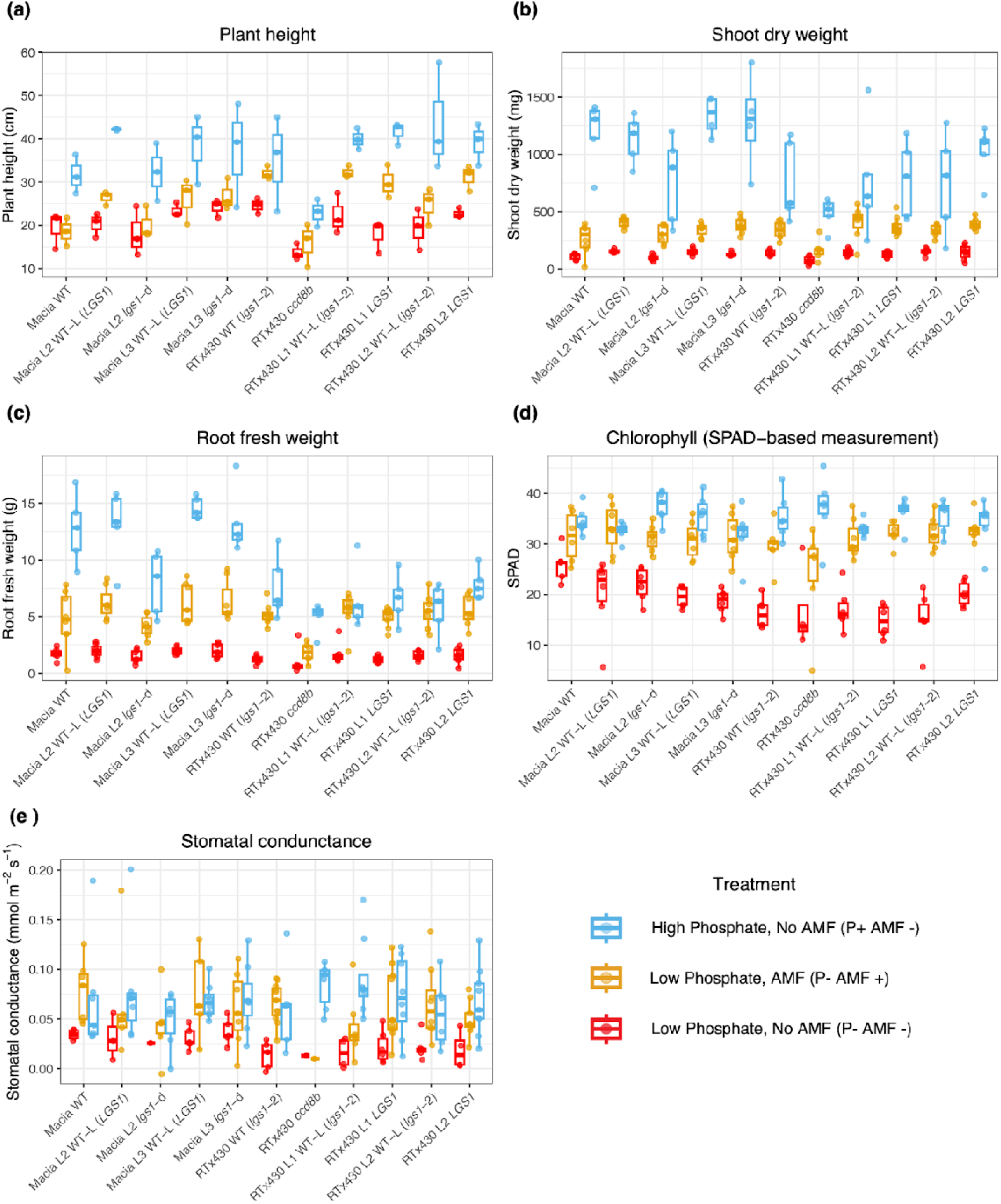
Phenotypes collected in greenhouse experiment. A) Plant height B) shoot dry weight C) root fresh weight of plants collected at Timepoint 2 (42 days). D) Single-photon avalanche diode (SPAD)-based measurement of leaf chlorophyll and E) Stomatal conductance collected at 28 days after plant emergence. Boxplots colored by treatment: high phosphate and no arbuscular mycorrhizal fungi (P+ AMF-; blue), low phosphate and arbuscular mycorrhizal fungi (P-AMF+ ; orange), low phosphate and no arbuscular mycorrhizal fungi (P-AMF-; red). Description of genotypes are reported in Table 1.

In all three treatments, the RTx430 *ccd8b* mutant was significantly shorter (Figure 4F, Figure 5A), had lower shoot weight (Figure 5B), and lower root weight (Figure 5C) as compared to the RTx430 WT. RTx430 *ccd8b* plants grown in the treatment without nutrient limitation (P+ AMF-) displayed phenotypic differences classically observed in strigolactone biosynthetic mutants. RTx430 *ccd8b* had increased shoot branching and tillering later in growth (data not shown), though this phenotype was not observed in RTx430 *ccd8b* plants grown in either low phosphate treatments with (P-AMF+) or without (P-AMF-) mycorrhizae, which may be attributable to a reduced growth rate under stressful conditions. These observations in the RTx430 *ccd8b* mutant support that Sb*CCD8b* loss-of-function results in pleiotropic effects, independent of nutrient availability. Nevertheless, under nutrient limitation, phenotypic effects were especially pronounced and growth was severely stunted compared to the RTx430 wild type (Figure 4E,F).

There were no significant differences in root or shoot weight for Macia *lgs1*-d grown in the high phosphate (P+ AMF-) treatment earlier in growth (21 days); however, later in growth (42 days) dry root biomass was significantly lower ( ^2^(1) = 4.18, *P* = 0.03, likelihood ratio test for fixed effect of *LGS1* deletion; Figure S10) as compared to the Macia WT and WT-L null segregants. As such, for plants of the Macia background differing in functionality of *LGS1* grown without nutrient limitation, differences in growth may be subtle, only detectable at later points in development, and *lgs1-*d plants may benefit less from the application of phosphate. For plants differing for the functionality of *LGS1* in the RTx430 background grown in the high phosphate treatment (P+ AMF-), we did not find evidence for any of the traits we measured (root weight, shoot weight, SPAD, or plant height) to be significantly different, again supporting that *LGS1* effects may depend on variation in other genes (Q4).

### Strigolactone biosynthesis deletion mutants have impaired colonization of mycorrhizae

At the first time point (21 days), RTx430 *ccd8b* plants had significantly delayed and overall reduced mycorrhizal colonization (t(9) = -6.65, *P* < 0.001, difference of genotypes by Student’s t-test; Figure 6B, Figure 7C), implicating Sb*CCD8b* to be involved in the early establishment of mycorrhizal partners. Furthermore, across both timepoints (21 and 42 days) RTx430 *ccd8b* root weight, shoot weight, and plant height remained significantly reduced as compared to RTx430 WT plants grown in the same treatment (Figure 5, Figure 6D, Figure 7C-D), indicating a relatively low benefit from the addition of mycorrhizae for plants with loss of function at Sb*CCD8b*. For RTx430 genotypes grown with (P-AMF+) and without (P-AMF-) mycorrhizae at 42 days, the relative increase in shoot weight was 20% lower for RTx430 *ccd8b* (t(8.2) = -4.53, *P* = 0.002, difference of genotypes by Student’s t-test; Figure 6D), suggesting that the impaired establishment of AMF, in turn, influenced the benefit mycorrhizal symbionts provided to RTx430 *ccd8b*.

**Figure 6.**
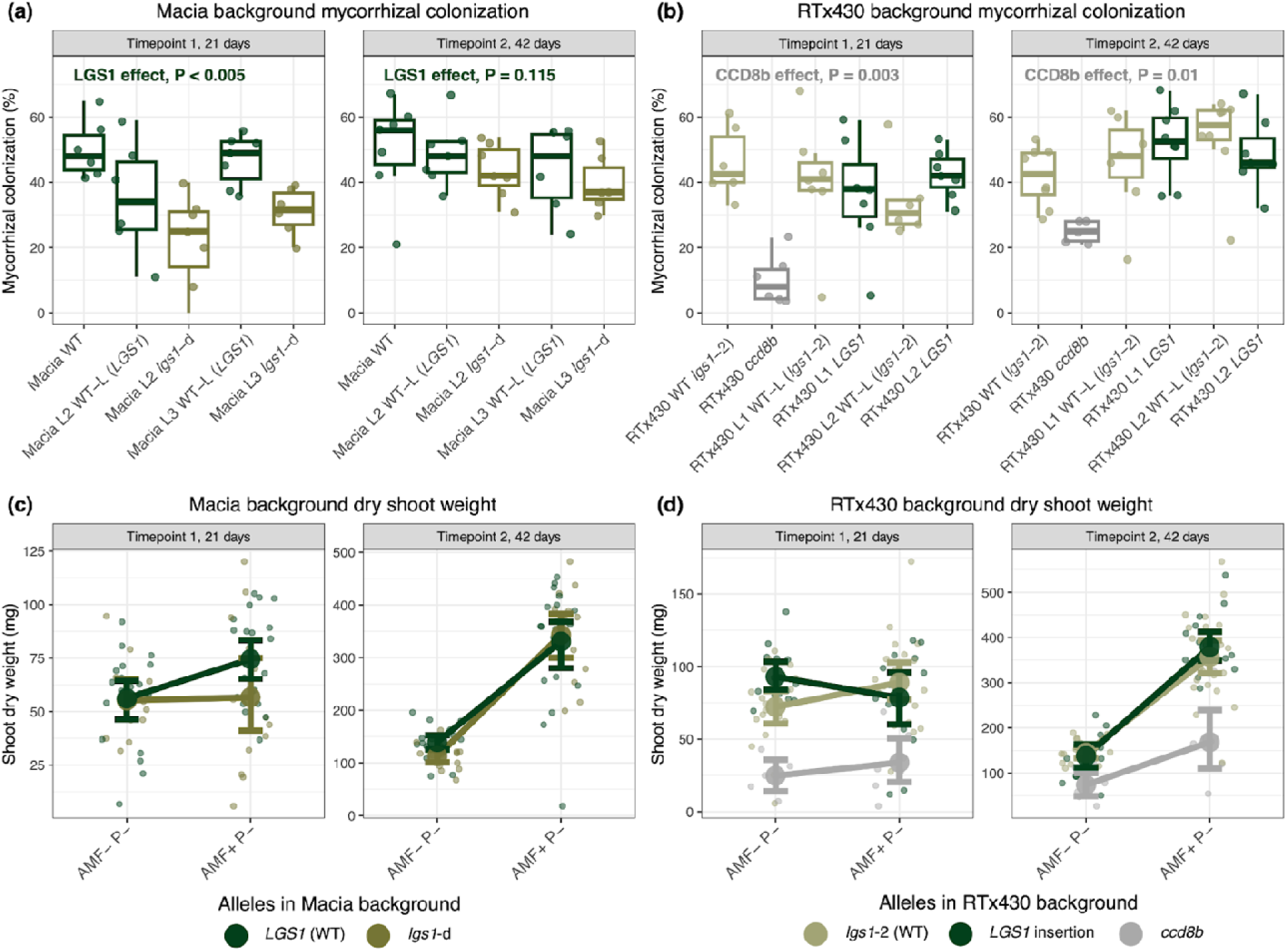
Mycorrhizal colonization is disrupted when strigolactone biosynthesis genes are knocked out of normally functional backgrounds. A) Percent of roots colonized by the mycorrhizal species *R. intraradices* at Timepoint 1 (21 days) and Timepoint 2 (42 days) for WT, “wild type-like” null segregants (WT-L), and *LGS1* deletion (*lgs1*-d) lines of the Macia background. B) Percent of roots colonized by mycorrhizal species *R. intraradices* at Timepoint 1 (21 days) and Timepoint 2 (42 days) for WT, *ccd8b* mutants, “wild type-like” null segregants (WT-L), and *LGS1* insertion mutants (*LGS1*) of the RTx430 background. WT-L and mutants of the same transformation event (L1,2,3) indicate paired genotypes. Significance of the allele effect (*LGS1* or Sb*CCD8b*, respectively) from linear models is reported. Dry shoot weight at Timepoint 1 (21 days) and Timepoint 2 (42 days) for C) Macia genotypes and D) RTx430 genotypes grown under low phosphate conditions, with (AMF+ P-) and without mycorrhizae (AMF-P-). The mean dry shoot weight for genotypes with the same allele grown in a treatment is plotted as a large dot with whiskers extending to the 95% confidence intervals.

**Figure 7.**
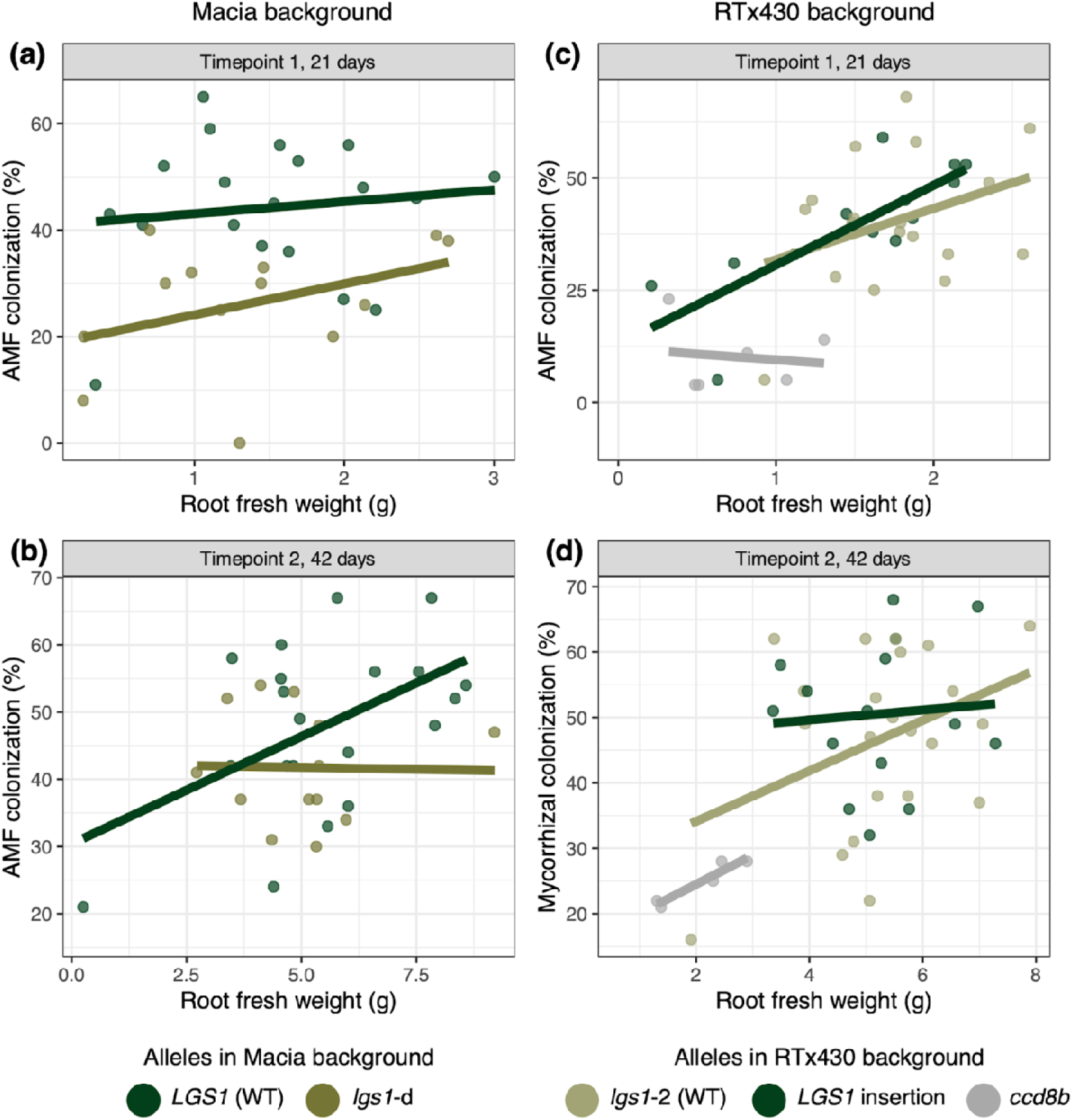
Percent roots colonized by mycorrhizae plot against root fresh weight. Genotypes of the Macia background at A) Timepoint 1 (21 days) and B) Timepoint 2 (42 days). Genotypes of the RTx430 background at C) Timepoint 1 (21 days) and D) Timepoint 2 (42 days). Points and regression lines are colored by allele.

For plants of the Macia background at the earlier time point (21 days), mycorrhizal colonization was on average 48% lower in Macia *lgs1*-d as compared to the colonization of WT and WT-L null segregants ( ^2^(1) = 11.38, *P <* 0.001, likelihood ratio test for fixed effect of *LGS1* deletion; Figure 6A, Figure 7A). This disruption in Macia *lgs1*-d early mycorrhizal colonization was also apparent from differences in shoot traits. At the earlier harvest time point (21 days) of the low phosphate mycorrhizal treatment (P-AMF+), Macia with functional *LGS1* had greater shoot weight than the Macia *lgs1*-d mutants (t(20) = -1.7327, *P* = 0.1, difference of genotypes by Student’s t-test; Figure 6C, Figure 4A-B). Additionally, plants of the Macia background with functional *LGS1* gained a greater benefit from the addition of mycorrhizae (t(38) = -2.69, *P* = 0.01, Macia *LGS1* difference between low phosphate mycorrhizal and non-mycorrhizal treatment by Student’s t-test), where Macia *lgs1*-d did not show any difference in average shoot weight between mycorrhizal (P-AMF+) and non-mycorrhizal (P-AMF-) treatments (Figure 6C).

At the later time point (42 days), while mycorrhizal colonization remained 15% lower for Macia *lgs1*-d as compared to colonization of WT and WT-L null segregants, the difference in colonization was no longer significant ( ^2^(1) = 2.48, *P* = 0.115, likelihood ratio test for fixed effect of *LGS1* deletion; Figure 6A,C, Figure 7B). Shoot dry weight between Macia *lgs1*-d and Macia lines with functional *LGS1* (WT, WT-L null segregants) at the later time point (42 days) increased in a similar manner for plants grown with (P-AMF+) and without (P-AMF-) mycorrhizae. This indicates that all Macia genotypes, regardless of *LGS1* functionality, benefit from the addition of mycorrhizae given sufficient time to associate with mycorrhizal symbionts. For plants of the RTx430 background, we did not find evidence for the functionality of *LGS1* to be associated with differences in mycorrhizal colonization. Across timepoints, mycorrhizal colonization was not significantly different between the RTx430 WT or WT-L plants maintaining the *lgs1*-2 deletion allele and the mutants with functional *LGS1* inserted (Figure 6B,D, Figure 7C,D).

## Discussion

Strigolactones are hormones that regulate plant development and physiology, while also acting as rhizosphere signals that mediate interactions with symbiotic microbes and parasitic plants. Our study explored the impact of strigolactones in sorghum, a crop vital to semiarid regions with often nutrient-poor soils, by examining mutants of the strigolactone biosynthesis genes Sb*CCD8b* and *LGS1*.

Artificial CRISPR-Cas9 knockouts of strigolactone biosynthesis genes affected many traits. Loss-of-function in Sb*CCD8b* led to changes in growth (Figure 5E,F), altered physiology (Figure 2D), disrupted mycorrhizal establishment (Figure 6B,D), significant changes to low-abundance root metabolites (Figure S7A), and changes in root architecture and anatomy (Figure 3). Similarly, deletion of functional *LGS1* in the Macia background (Macia *lgs1*-d) resulted in transcriptional changes of shoots enriched for promoter motifs for stress response and growth pathways (Table S2), a down-regulation of root metabolites and up-regulation of shoot metabolites (Figure S7C-H), reduced carbon assimilation (Figure 2A), impaired early mycorrhizal colonization (Figure 6A,C), and reductions in root growth under non-limiting conditions (Figure S10). Together, our study demonstrates that deletion of the tested strigolactone biosynthesis genes impacts both endogenous developmental processes and exogenous signaling functions, leading to measurable trade-offs in plant performance (Q3). At the same time, we hypothesize epistasis with a neighboring strigolactone biosynthesis gene may modify the effects of loss-of-function at *LGS1*.

### Genetic background dependent trade-offs and evidence for epistasis

A key finding of our study is that the magnitude and direction of *LGS1*-associated trade-offs depend strongly on genetic background (Q4). While deletion of *LGS1* in the Macia background resulted in clear impeded photosynthesis and AMF colonization, reintroduction of functional *LGS1* into the RTx430 background restored parasite susceptibility (Figure 1) without enhancing photosynthesis or AMF colonization. Aside from increased nodal root number under high phosphate conditions (Figure S9B), RTx430 mutants with functional *LGS1* inserted showed minimal evidence for differences compared to RTx430 WT and WT-L null segregants (*lgs1*-2). This apparent epistasis may be due to another gene near *LGS1* involved in strigolactone biosynthesis. In addition to *LGS1* (*Sobic.005G213600*), the *lgs1*-2 deletion allele carried by RTx430 also includes deletion of *Sobic.005G213400* and *Sobic.005G213500* (*Sb3500*) (Table 2), genes we did not insert into RTx430 along with *LGS1*. *Sb3500* is a 2-oxoglutarate-dependent dioxygenase affecting the conversion of carlactone to 5DS (Yoda et al., 2023) and is ∼3 kb upstream of the *LGS1* gene. Yoda et al. 2023 previously showed epistasis between *Sb3500* and *LGS1* for strigolactone synthesis when expressed with other strigolactone biosynthetic genes in *Nicotiana benthamiana* (expected strigolactones shown in Table 2), suggesting the insertion mutant RTx430 *LGS1* did not have the typical strigolactone profile of Macia WT. It may be that restoration of functional *Sb3500* and a primarily 5DS strigolactone profile would be required to enhance photosynthesis and AMF colonization of RTx430. Though we found only minimal phenotypic differences between RTx430 WT and the RTx430 *LGS1* insertion mutant, the combination of alles in the *LGS1* insertion mutant, absent *Sb3500* and present *LGS1*, is extremely rare in natural sorghum genotypes (only 1 of 704 landraces, Table 2) and may indicate fitness costs associated with this allelic combination. RTx430 is an extensively used breeding line (Gurel et al., 2009) for both research and hybrid production, and other natural mutations in RTx430 may influence *LGS1* deletion effects. Additionally, recent studies suggest some endogenous strigolactone signalling is driven by non-canonical strigolactones and not canonical (e.g. 5DS, orobanchol) strigolactones (Ito et al., 2022), and so the effects of mutations in canonical strigolactone biosynthesis genes like *LGS1* and resulting feedbacks require more dissection. Future work would benefit from disentangling the allelic effects of *LGS1* and *Sb3500* from differences in genetic background, and functionally validating the impact of *Sb3500* on sorghum strigolactone exudate profiles.

### Linking strigolactones, stress responses, and differences in photosynthesis

The pleiotropic effects observed in the strigolactone biosynthesis mutants used in experiments are consistent with known crosstalk between strigolactones and stress-response pathways (Q2). Strigolactones interact with abscisic acid (ABA) signaling and contribute to plant responses to drought and other abiotic stresses (Korek and Marzec, 2023). In our study, Sb*CCD8b* mutants in the RTx430 background trended towards elevated stomatal conductance earlier in growth (Figure S3C) and RTx430 *ccd8b* roots had an increased accumulation of dihydroorotate (Table S3), a metabolite associated with drought-sensitive genotypes (Li et al., 2022). Additionally, both Sb*CCD8b* and *LGS1* knockout mutants exhibited reduced net carbon assimilation (Figure 2A,C; Figure S4A). In Macia *lgs1*-d, this reduction was related to decreases in maximum rate of carboxylation (*V*_cmax_) (Figure S4D), suggesting limitations to Rubisco activity when functional *LGS1* is deleted. Relatedly, enrichment of AP2/EREBP transcription factor binding sites in the promoter regions of Macia *LGS1* knockouts (Macia *lgs1*-d) differentially expressed genes supports a regulatory basis for these effects. This family of transcription factors is involved in both stress responses (Xu et al., 2011) and photosynthetic regulation (Hong et al., 2009). As such, AP2/EREBPs have been suggested as ideal candidates for crop improvement and their overexpression enhances tolerances to multiple abiotic and biotic stressors (Xu et al., 2011). For example, overexpression of the At*DREB1A* (AP2/EREBP subfamily transcription factor) in Chrysanthemum resulted in significantly higher photosynthetic capacity, and elevated activity of Rubisco (Hong et al., 2009). Together, the decreased carbon assimilation rate in Macia *lgs1*-d may be attributable to the AP2/EREP or other associated regulatory networks, although additional work is needed to resolve the underlying mechanisms.

### Developmental regulation and the miR156–SPL pathway

Many of the phenotypes altered in the strigolactone biosynthesis knockout mutants, including - reduced stomatal density, decreased mycorrhizal colonization, altered photosynthetic rates, and increased nodal root production - are known to be regulated by the miR156-SPL pathway (Poethig, 2013; Aung et al., 2015; Lawrence et al., 2021) which regulates developmental timing and the transition of vegetative characteristics from juvenile to adult (Q2). The majority of phenotypes we collected in greenhouse experiments were collected from plants transitioning from the juvenile to the adult stage (average last juvenile leaf = node 3, average first adult leaf = node 7, all leaves between these positions are considered transition leaves) and traits observed in the RTx430 *ccd8b* and Macia *lgs1*-d mutants are consistent with a more “juvenile-like” developmental state, associated with elevated miR156 and reduced SPL expression. Previous studies have linked the miR156-SPL pathway with strigolactone biosynthesis and signaling. SPL proteins mediate strigolactone signaling through direct interaction with the signaling molecule DWARF53 (D53), which binds SPLs in the absence of strigolactones suppressing their transcriptional activity (Barbier et al., 2023). Further, SPLs promote expression of *CCD* and *NCED* genes, key enzymes in strigolactone biosynthesis (Arango et al., 2016). Supporting this, the miR156-SPL pathway regulates miR172 expression, which targets AP2 transcription factors (Ó’Maoiléidigh et al., 2021), the largest group of transcription factors enriched in our analysis of regulatory pathways related to patterns of differential expression in Macia *lgs1*-d mutants (Table S2). Consequently, the range of phenotypes altered when functional strigolactone biosynthesis genes are knocked out (RTx430 *ccd8b* and Macia *lgs1*-d) may be associated with, or potentially mediated by, changes in developmental timing through the miR156-SPL regulatory network.

### Impacts on nutrient signaling and mycorrhizal symbiosis

Strigolactones play a central role in nutrient signaling, particularly under phosphate limitation, where they promote root architectural changes and facilitate recruitment of mycorrhizal symbionts (Mayzlish-Gati et al., 2010; Ruyter-Spira et al., 2011; Alder et al., 2012; Sun et al., 2014). Previous observations by Hao et al. (2023) demonstrated significant shifts in the root-associated microbiome in Sb*CCD8b* knockouts. In agreement with these findings, we found the same RTx430 *ccd8b* knockout lines exhibited significantly reduced colonization by arbuscular mycorrhizal fungi and diminished benefits from this symbiotic partner (Q1, Figure 6B,D). We additionally demonstrate the disruption of early mycorrhizal colonization for Macia *lgs1*-d (Figure 6A, Figure 7A), though given sufficient time, the relative colonization and benefit of mycorrhizae appears similar for plants of the Macia background regardless of *LGS1* functionality (Q1, Figure 7B). The effects of impaired symbiosis with AMF may arise from multiple mechanisms, including altered strigolactone stereochemistry, reduced quantify strigolactone in root exudates, or both. Results from *S. hermonthica* germination assays reported in this study and previously (Bellis et al. 2020) are consistent with the hypothesis that absence of functional *LGS1* reduces total strigolactone exudation. Additionally, both orobanchol and 5DS are known to stimulate mycorrhizal branching (Akiyama et al., 2005; Akiyama et al., 2010). Reduced early colonization for Macia *lgs1*-d may therefore reflect insufficient signaling through reduced strigolactone exudation (Figure 7A), while the recovery at the later time point may indicate cumulative and sufficient signaling over time. In contrast, RTx430 *LGS1* insertion mutants showed no difference in mycorrhizal colonization at either time point (Figure 6B,D), despite a clear difference in strigolactone profiles from RTx430 WT and WT-L plants (*lgs1*-2), as evidenced by elevated germinability of parasite populations (Q1, Figure 1A-B). The *LGS1* insertion in the RTx430 background is expected to shift strigolactone composition from orobanchol toward 5DS, though likely to a lesser extent than in Macia WT due to the absence of *Sb3500* (Table 2). As plant responses to mycorrhizal colonization can vary across fungal species and strains (Angelard et al., 2010) and the same sorghum genotype can have substantial variation in colonization rates between fungal species (Gobena et al., 2017), our conclusions are limited to this *R. intraradices* experiment.

### Implications for breeding and deployment of LGS1 alleles

From an applied perspective, *LGS1* loss-of-function alleles are of considerable interest for improving sorghum’s resistance to *S. hermonthica* (Hess et al., 1992; Ejeta, 2007). Our results demonstrate that such alleles, specifically deleting functional *LGS1* from a normally functional background (Macia), reduce parasite germination (Bellis et al. 2020) but may also be coupled with trade-offs in growth, photosynthesis, and early establishment of mycorrhizal symbiosis. The absence of strong trade-offs in comparing the RTx430 WT (*lgs1*-2) to mutants with functional *LGS1* inserted suggests that these costs are not inevitable and may be mitigated by genetic background; however, the rarity of the combination of alleles present in the RTx430 *LGS1* insertion mutant may suggest other fitness trade-offs we did not observe. Epistasis, in particular genetic backgrounds or between multiple genes that impact sorghum strigolactone profiles (e.g. *Sb3500* and *LGS1*), may therefore represent vital variation for optimizing plant performance while maintaining resistance to *S. hermonthica* in farmer-preferred varieties. At the same time, we caution that the experiments described here likely do not fully capture the dynamics of field-grown sorghum, and any trade-offs incurred from deleting *LGS1* from a normally functional background could be preferable to crop loss caused by *S. hermonthica* pressure. Our study advances understanding of strigolactone biology in sorghum and highlights the complexity of co-evolutionary dynamics, where functional variation at a single locus can confer parasite resistance but with evidence for costs to host fitness.

## Acknowledgements & Funding

We thank Scott Diloreto and Sam Gruneberg for sharing their greenhouse and plant care expertise. We are grateful to Amanda Penn, Shiran Ben-Zeev, Diana Gamba, and Eric Nascari for their assistance with greenhouse data collection. We also thank the Corteva Sorghum transformation team for generation of RTx430 transgenic events, control environment team for taking care of and backcrossing plants, and the analysis team for molecular characterization of the material. RJHS is funded by the USDA National Institute of Food and Agriculture and Hatch Appropriations under Project No. PEN05039 and Accession No. 7008935, as well as by the USDA project “The Genetic Basis of Maize Response to Arbuscular Mycorrhizal Fungi” (Grant No. 2022-67013-38264). DPS acknowledges support from the USDA National Institute of Food and Agriculture (Grant No. 2020-67019-31796). JJK is supported by USDA NIFA and Hatch Appropriations under Project No. PEN04956 and Accession No. 7006496. JRL, JM, and CMM acknowledge support from the Bill & Melinda Gates Foundation.

## Author Contributions

CMM, JRL, RHJS conceived and planned the research. CMM, MT, JM, EJA, AC, AC, YX, EHL-P, and MP performed the experiments and analyzed data. Plant genetic material was developed by DPS and HG. *Striga hermonthica* seed was collected by SR and BN. CMM, JRL, and EHL-P wrote the paper with input from all authors.

