## SupplementaryMaterial for "Strigolactone effects on *Sorghum bicolor* ecophysiology and symbioses"

**PDF includes the following supplementary materials**

Supplementary Methods

Supplementary Figures S1 - S11

Supplementary Tables S1 - S6

References for Supplementary Methods

**Supplementary Methods**

*Backcross sorghum LGS1 deletion variants to wild type Macia*

Hot-water emasculation was applied when backcrossing the *LGS1* deletion variants ($S_2$ transgenics, Macia *lgs1*-d; Figure S1) to wild type Macia. The method was developed by Stephens and Quinby in 1934 and has been used routinely (Schertz and Dalton 1980) with minor modification here. A panicle that has just begun to flower is chosen. Any branch that had pollen shed along with immature branches at the lower part of the panicle were removed. The panicle was then submerged in a hot water bath at 45C for ~13 minutes followed by 1-3 days drying. The dried panicles were crossed by tying with *LGS1* deletion variant ($S_2$ plants) panicles together or carrying pollen from *LGS1* deletion variant panicles to produce $BC_0$ seeds.

To verify if crossing was successful, PCR was performed to detect *LGS1* deletion in the $BC_0$ plants as described previously (Bellis et al. 2020). $BC_0$ plants positive with *LGS1* deletion were selfed to make $BC_0F_2$ seeds, these $BC_0F_2$ seeds segregated at 1:2:1 (homozygous: hemizygous: null *LGS1* gene deletion), homozygous (*lgs1*-d) and null *LGS1* deletion (*LGS1* present) plants produced more seeds for further evaluation.

*RTx430* LGS1 *transgenic event generation*

*LGS1* gene is not present in the RTx430 genetic background. To further demonstrate *LGS1* gene in relation to *Striga* tolerance, we introduced the functional *LGS1* gene to the RTx430 genetic background via *agrobacterium* random transformation.

*Agrobacterium tumefaciens* strain and plasmid vectors

The *LGS1* gene (*Sobic.005G213600*), a 4070bp genomic sequence including a 2kb sequence served as native promoter, 5' UTR, exon 1, intron 1, exon 2, and a 530bp of 3 UTR sequence was cloned into the binary vector pPHP98108 using standard cloning technology. To achieve marker-free in transformation process, the excision cassette in the binary vector contains *Wus2* ($Pltp_{pro}$:*Wus2*), a heat shock inducible *moCRE* gene ($Hsp17.7_{pro}$:*moCRE*) and a selectable marker ($Zm\text{-}Ubi_{pro}$:*NPTII*) flanked by the directly repeated *lox*P sites (Figure S11). The binary vector was electroporated into *Agrobacterium* strain LBA4404 Thy- containing the accessory plasmid pPHP71539, recombinant colonies were selected on media supplemented with both gentamicin and spectinomycin. The construct was then subjected to next-generation sequencing and sequence confirmation before transformation experiments, for further detail, see Che et al. (2022).

*Plant material, sorghum transformation, and transgenic event quality analysis*

Immature embryos isolated from sorghum plants were transformed with *Agrobacterium* auxotrophic strain LBA4404 Thy- carrying a ternary vector transformation system to generate transgenic sorghum plants (Anand 2018; Wu 2018). The *Pltp:WUS2* gene enables sorghum transformation and eliminates the nine-week callus proliferation phase. Following the *Agrobacterium* infection and co-cultivation steps, the immature embryos were sub-cultured on the multi-purpose medium with selection for three weeks to induce somatic embryo formation.

After three weeks of stringent selection, Wus2/moCRE/NPTII cassette excision was carried out by heat-shock treatment at 45 °C for 3 h (Che et al 2022). Embryos were transferred to the maturation medium without selection for four weeks then placed on rooting plates as described in Anand and Wu (2018). $T_0$ plants were planted in a soil flat and grown in a controlled greenhouse environment.

$T_0$ plants were subject to molecular analysis. Leaf DNA was extracted as described previously (Bellis et al. 2020; Che et al. 2022). Quantity PCRs were performed as described in Che et al. (2022) with primers listed in Table S6. The $T_0$ plants with successful marker excision (Figure S11) and only one copy of the *LGS1* gene insertion were further analyzed by Southern by sequencing (Gina et al. 2015) to confirm the *LGS1* gene insertion and excision success. Selected $T_0$ plants were used to produce $T_1$ and later progeny seeds. SbS analysis showed *LGS1* gene (Figure S11) were inserted on Chr01 for the L1 transformation event and Chr02 for the L2 transformation event. Homozygous and event null (WT-L) $T_2$ seeds from the 2 selected *LGS1* insertion lines were collected and evaluated in this study.

#### Supplementary Figures

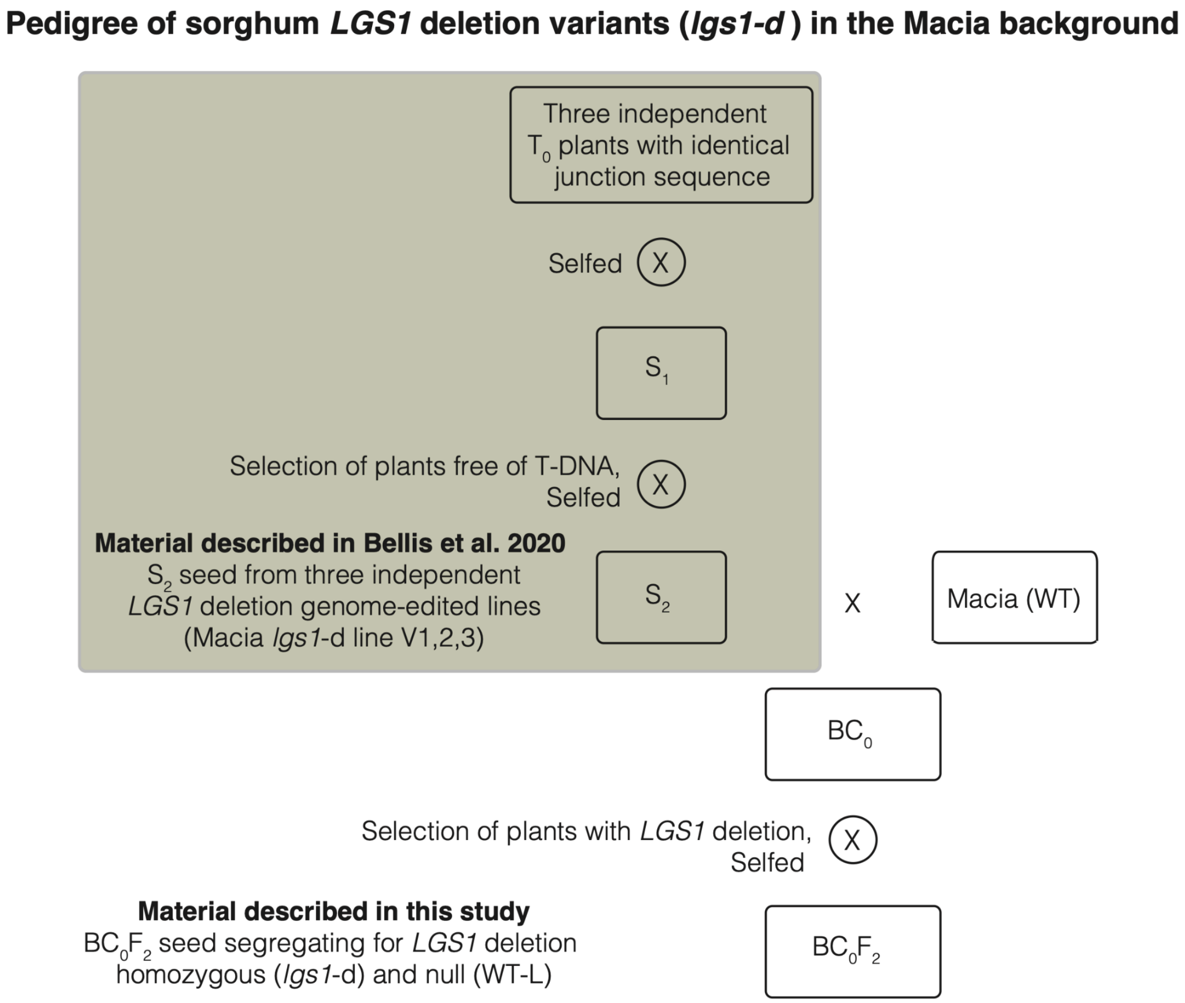


**Figure S1**

Pedigree of *lgs1*-d mutants generated in the Macia background. The green box is the crossing scheme for material of the $S_2$ generation as described in Bellis et al. (2020). The material described in this study is $BC_0F_2$ seed. $BC_0F_2$ seed was generated from crossing $S_2$ seed to Macia WT, additional rounds of selfing, and selection of seed segregating for homozygous deletion (*lgs1*-d) and “wild type-like” null event mutants (WT-L).

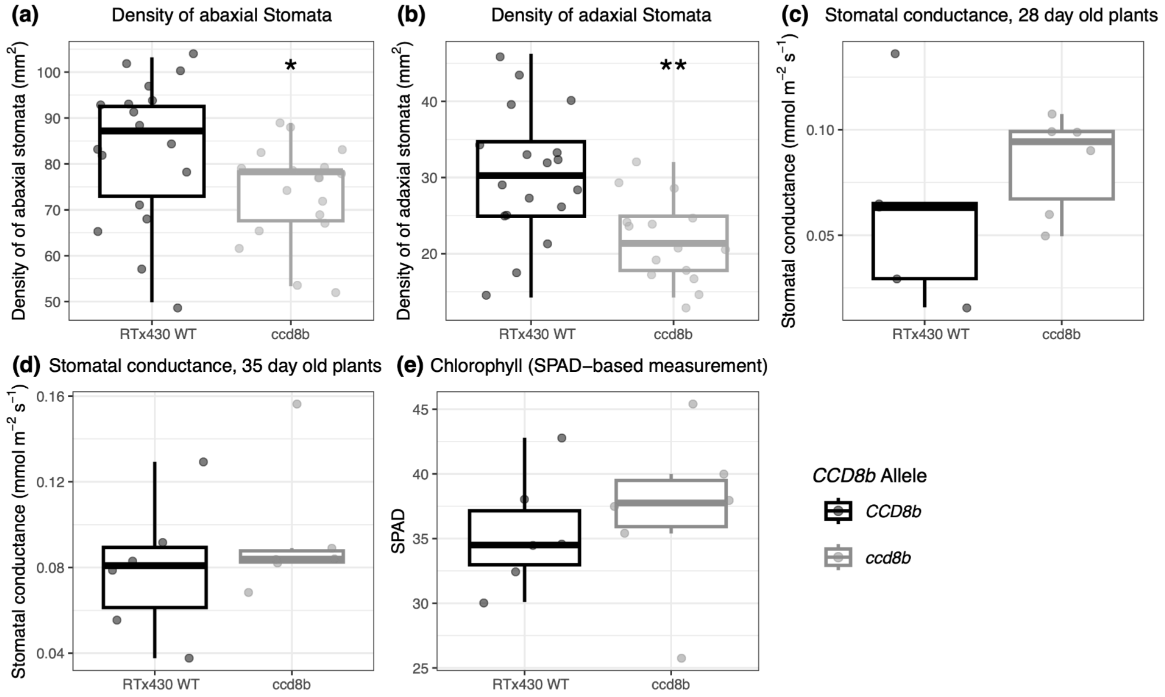


**Figure S2**

Stomatal and chlorophyll traits comparing *ccd8b* mutants and WT in the RTx430 background. Plants grown without nutrient limitation. A) Density of abaxial stomata. B) Density of adaxial stomata. C) Stomatal conductance measured with 28 day old plants, data same as shown in Figure 5E. D) Stomatal conductance measured with 35 day old plants. E) SPAD-based measurement of chlorophyll content measured with 28 day old plants, data same as shown in Figure 5D. Asterisks denote a significant *P*-value comparing WT and mutant by Student's *t*-test. *$P<0.05$, **$P<0.01$

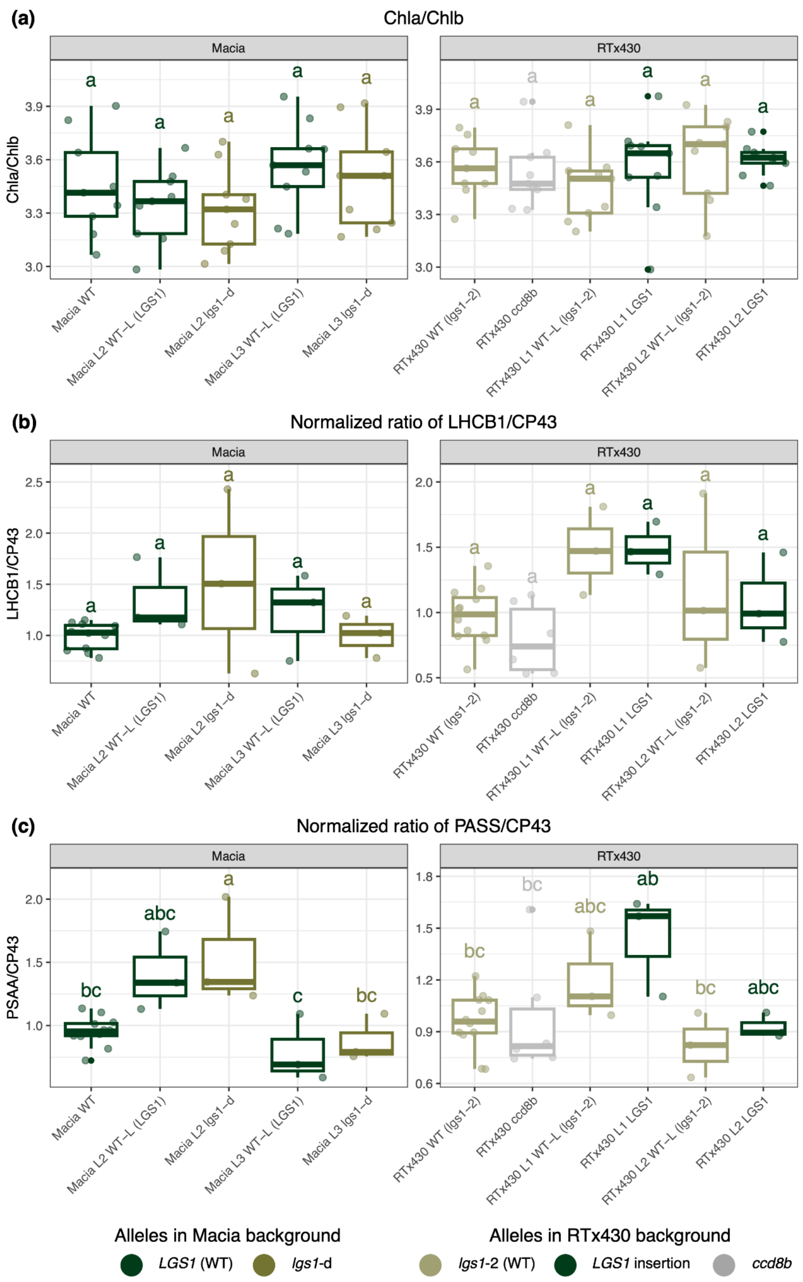


**Figure S3**

A) Ratio of Chl*a*/Chl*b*. B) Ratio of PSAA/CP43. C) Ratio of LHCB1/CP43. Protein ratios are normalized against a genotype's corresponding wild type (Macia WT or RTx430 WT). Genotypes of the Macia background include the WT, "wild type-like" null segregants (WT-L), and *LGS1* deletion mutants (*lgs1*-d). Genotypes of the RTx430 background include the WT, *ccd8b* mutant, "wild type-like" null segregants (WT-L), and *LGS1* insertion mutants (*LGS1*). WT-L and mutants of the same transformation event (L1,2,3) indicate paired genotypes. Genotypes that share the same letter are not significantly different as determined by Tukey's post hoc test.

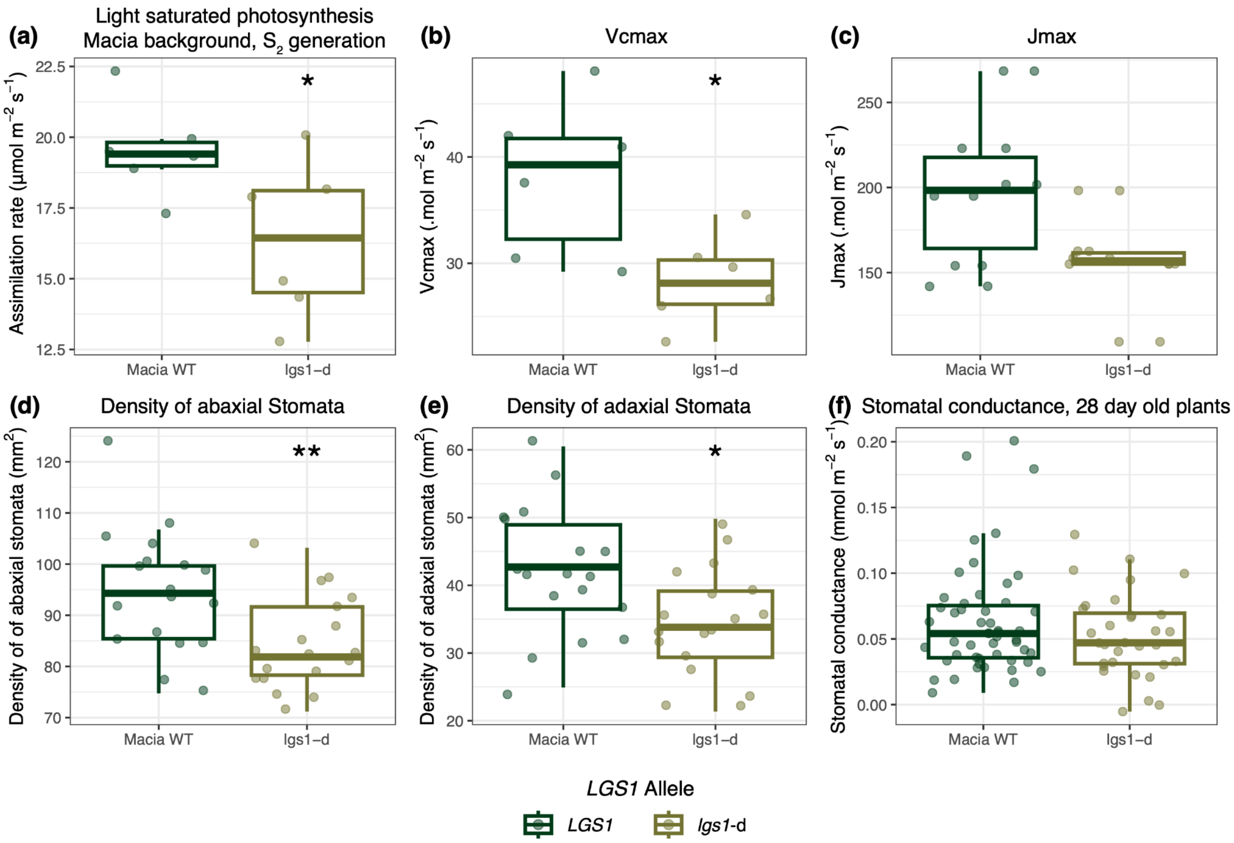


**Figure S4**

Physiological traits, traits estimated from $A_{\text{net}}$–$C_i$ curves, and stomatal traits comparing *LGS1* deletion (*lgs1*-d) mutants to the WT in the Macia background. Plants grown without nutrient limitation and all data measured 28 day old plants. A) Net carbon assimilation rate using Macia *lgs1*-d mutants of the $S_2$ generation. B) $V_{\text{cmax}}$ estimated from $A_{\text{net}}$–$C_i$ curve shown in Figure 2B. C) $J_{\text{max}}$ estimated from $A_{\text{net}}$–$C_i$ curves shown in Figure 2B. D) Density of abaxial stomata. E) Density of adaxial stomata. F) Stomatal conductance, data same as shown in Figure 5E. Asterisks denote a significant $P$-value comparing WT and mutant by Student's $t$-test. *$P$<0.05, **$P$<0.01

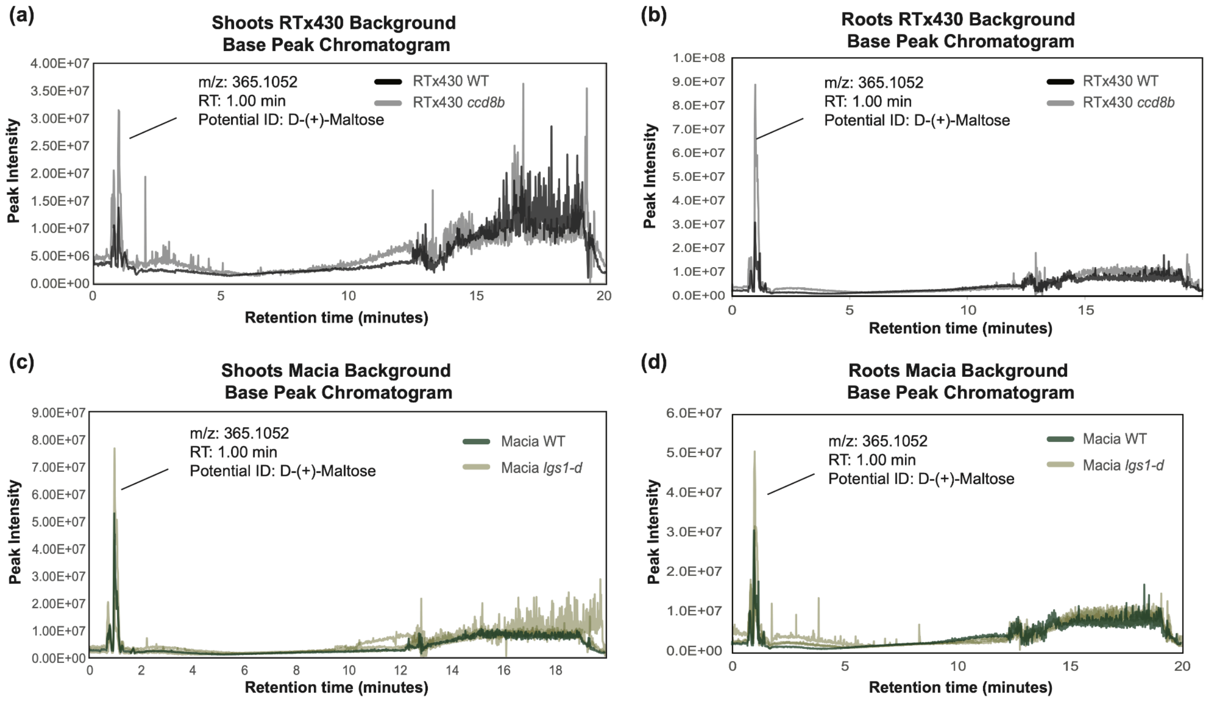


**Figure S5**

Base peak chromatograms (BPCs) for RTx430 WT (black) and *ccd8b* mutant (gray) A) shoots and B) roots. BPCs for Macia WT (dark green) and three *LGS1* deletion mutants of the $S_2$ generation from three separate transformation events (*lgs1*-d, light green overlaid with high opacity) C) shoots and D) roots.

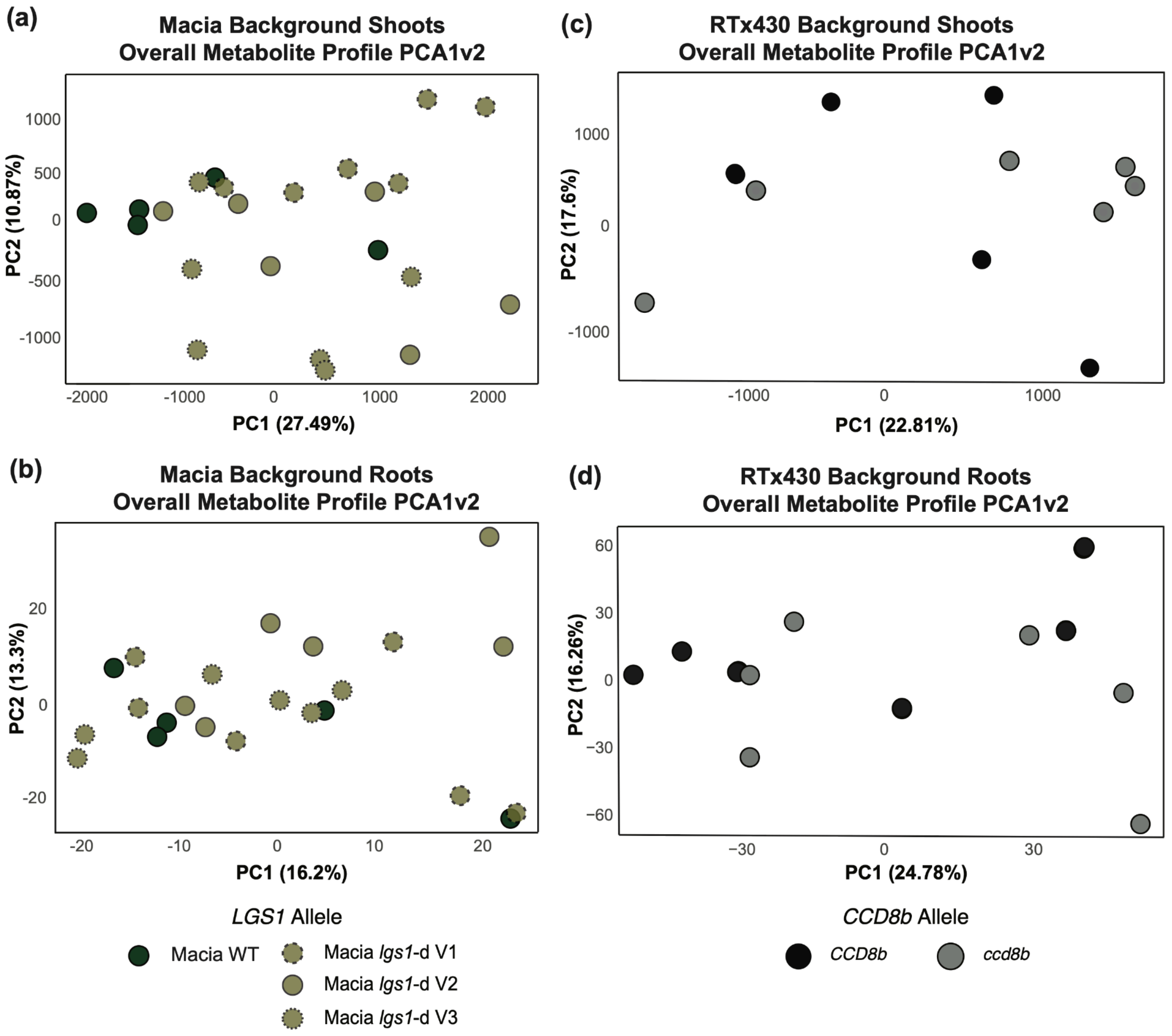


**Figure S6**

The first two principal component axes (PC1, PC2) of overall metabolite profiles for Macia WT (dark green) and three *LGS1* deletion mutants (*lgs1*-d, light green) of the $S_2$ generation from three separate transformation events (V1,2,3) A) shoots and B) roots. PCA of overall metabolite profiles for RTx430 WT (black) and *ccd8b* mutant (gray) C) shoots and D) roots.

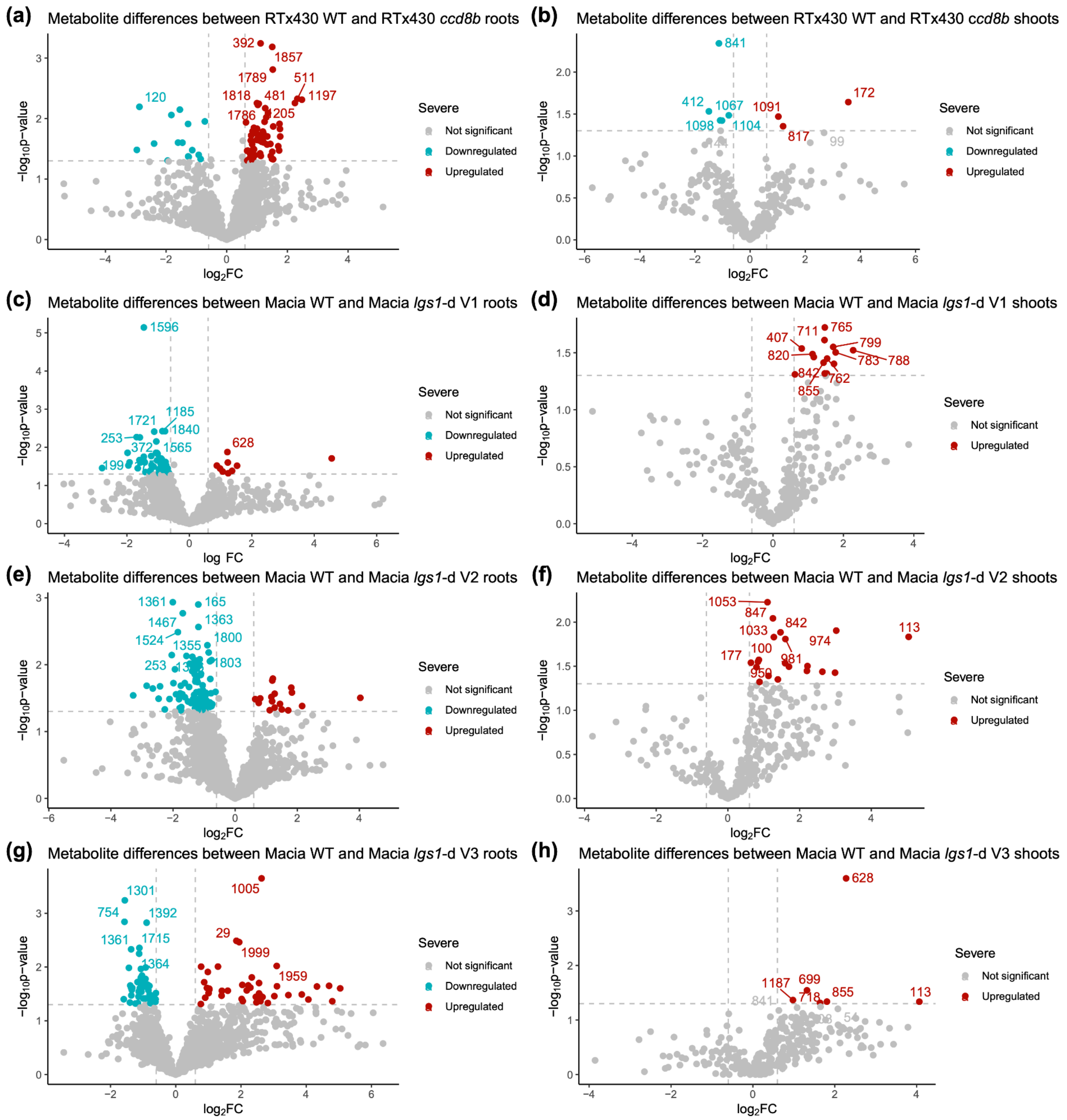


#### Figure S7

Volcano plot of differentially accumulated metabolites comparing the RTx430 WT and *ccd8b* mutant A) roots and B) shoots. Volcano plot of differentially accumulated metabolites comparing the Macia WT and *LGS1* deletion lines of the $S_2$ generation from three separate transformation events (*lgs1*-d V1,2,3) C, E, G) roots and D, F, H) shoots. Significantly up-regulated metabolites are colored in red and significantly down-regulated metabolites are in blue. Annotation information on significantly accumulated metabolites can be found in Table S3, S4.

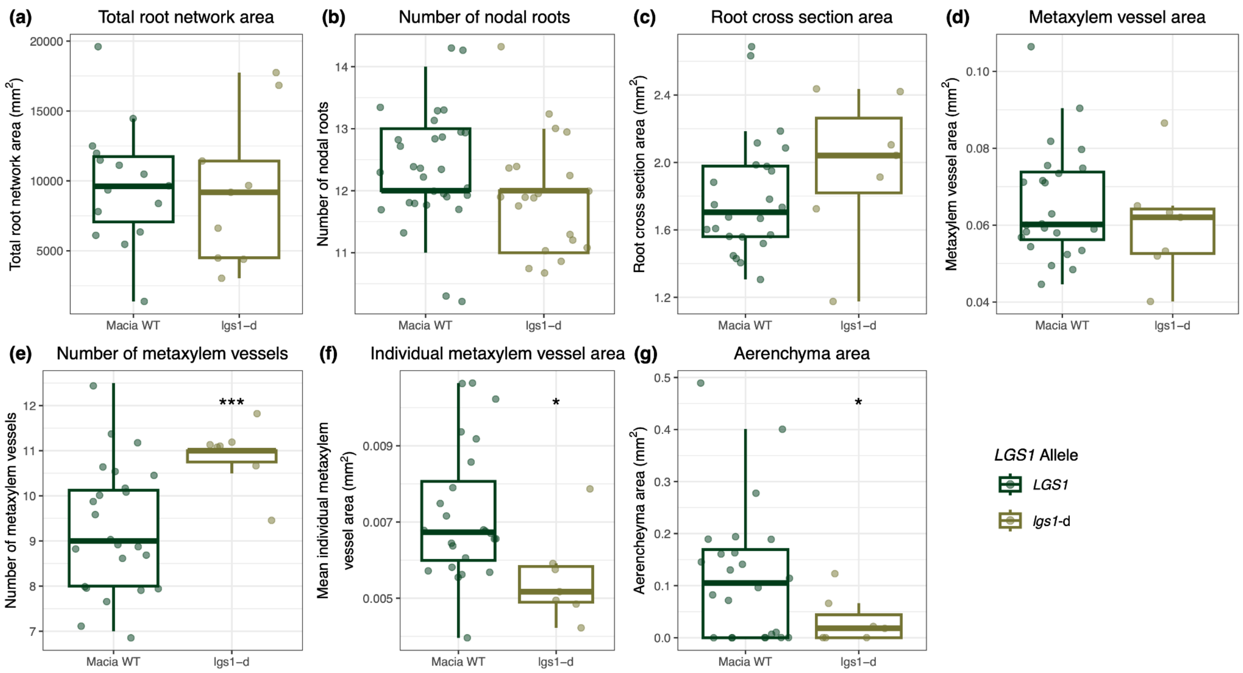


### Figure S8

Root Architecture and anatomy for *LGS1* deletion (*lgs1*-d) mutants as compared to the WT in the Macia background. Points plotted for "Macia WT" includes the untransformed wild type and "wild type-like" null segregants (WT-L) of two transformation events, and for "*lgs1*-d" includes mutants of two transformation events. A) Total root system network area. B) Number of nodal roots. C) Root cross-section area. D) Total metaxylem vessel area. E) Number of metaxylem vessels. F) Individual metaxylem vessel area. G) Aerenchyma area. Asterisks denote a significant *P*-value comparing WT and mutant by Student's *t*-test. *$P<0.05$, ***$P<0.0001$

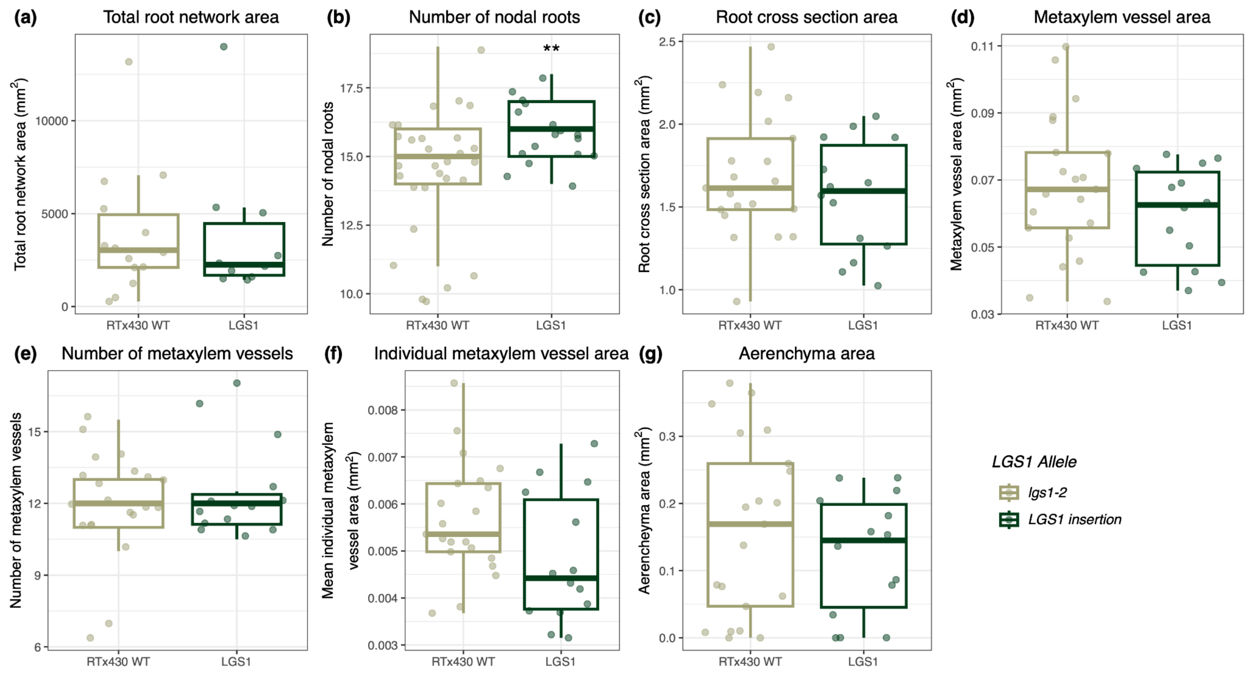


**Figure S9**

Root Architecture and anatomy for *LGS1* insertion (*LGS1*) mutants as compared to the WT (*lgs1*-2) in the RTx430 background. Points plotted for "RTx430 WT" includes the untransformed wild type and "wild type-like" null segregants (WT-L) of two transformation events, and for "*LGS1*" includes mutants of two transformation events. A) Total root system network area. B) Number of nodal roots. C) Root cross-section area. D) Total metaxylem vessel area. E) Number of metaxylem vessels. F) Individual metaxylem vessel area. G) Aerenchyma area. Asterisks denote a significant *P*-value comparing WT and mutant by Student's *t*-test. $^{**}P<0.01$

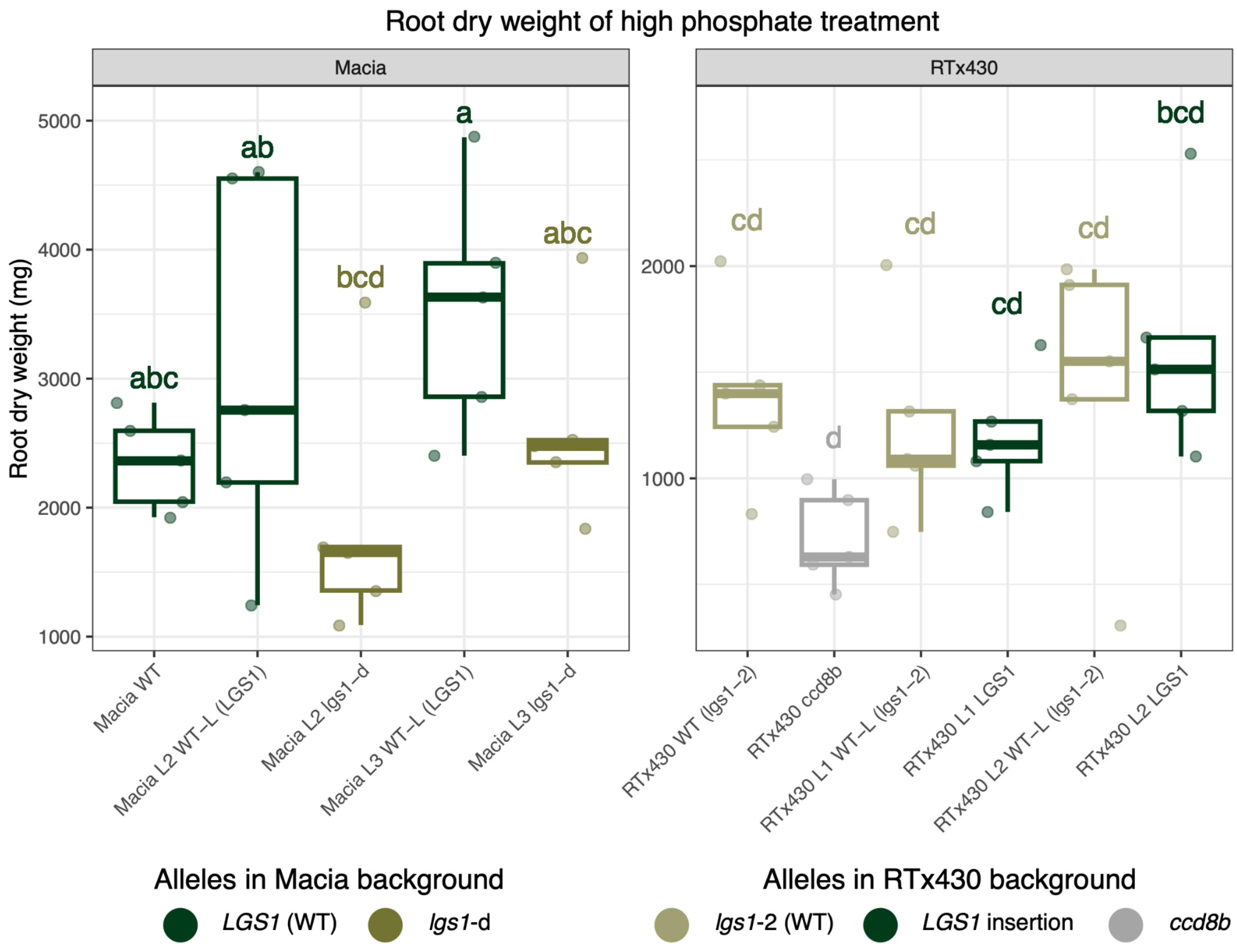


**Figure S10**

Root dry weight of plants grown in the high phosphate, no mycorrhizae treatment (P+, AMF-) collected at the second timepoint (42 days). Genotypes of the Macia background include the WT, "wild type-like" null segregants (WT-L), and *LGS1* deletion (*lgs1*-d). Genotypes of the RTx430 background include the WT, *ccd8b* mutant, "wild type-like" null segregants (WT-L), and *LGS1* insertion mutants (*LGS1*). WT-L and mutants of the same transformation event (L1,2,3) indicate paired genotypes. Boxplots and points colored by allele. Genotypes that share the same letter are not significantly different as determined by Tukey's post hoc test. ($n$ = 5).

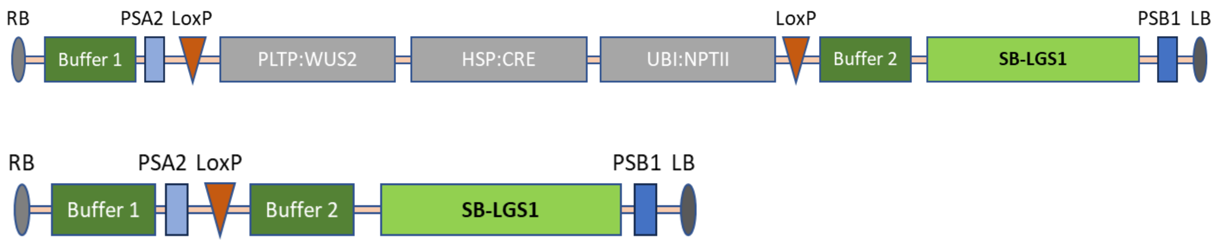


**Figure S11**

Insertion of the *LGS1* gene to RTx430 genome with random *agrobacterium* transformation. (Top) Schematic drawing of the T-DNA used to randomly introduce the *LGS1* genomic sequence to the RTx430 genome in the early transformation stage. Buffer 1 and Buffer 2 are sorghum gene's terminator sequences to prevent interference from other sequences to the *LGS1* gene expression, *WUS2* to enhance transformation, *NPTII* serves as transformation selection marker, and CRE to promote recombination of the two loxP, it is controlled by heat shock promoter. (Bottom) Remaining T-DNA in $T_0$ plants after heat shot treatment, heat shot before regeneration activated CRE gene to excise all sequences between the 2 loxP sites. PSA2 and PSB1 are unique sequences included in T-DNA for primer design.

**Table S1**

Known transcription factor binding sites (TFBS) overlapping the most frequently identified motif in promoter regions of Macia *LGS1* deletion mutants (*lgs1-d*) differentially expressed root genes. The discovered motif (GACCGTCCATC) shows no significant enrichment in differentially expressed genes relative to non-differentially expressed genes ($P = 0.48$). For each TFBS, the table reports the TFBS name (motif ID), motif name, TF class, *P*-value, and the number of overlapping base pairs between the discovered motif and the TFBS motif sequence.

| **Discovered Motif** | **TFBS name** | **Motif name** | **Class** | ***P*-value** | **Overlap (bp)** | **TFBS motif sequence** |
|---|---|---|---|---|---|---|
| GACCGTCCATC | MA1048.1 | ERF018 | AP2/EREBP | 0.000590171 | 8 | ACCGACCA |
| GACCGTCCATC | MA1234.1 | ERF017 | AP2/EREBP | 0.00101979 | 10 | CCTCCACCGTCCAT |
| GACCGTCCATC | MA1799.1 | SPL10 | C2H2 zinc finger factors | 0.00122548 | 10 | GTCCGTACAA |
| GACCGTCCATC | MA1795.1 | RAP2-4 | AP2/EREBP | 0.00192098 | 10 | CACCGACCAT |
| GACCGTCCATC | MA1057.1 | SPL12 | C2H2 zinc finger factors | 0.00375907 | 8 | ACCGTACA |
| GACCGTCCATC | MA1600.1 | ZNF684 | C2H2 zinc finger factors | 0.00424291 | 11 | ATACAGTCCACCCCTTTA |

**Table S2**

Known transcription factor binding sites (TFBS) overlapping the most frequently identified motif in promoter regions of Macia *LGS1* deletion mutants (*lgs1-d*) differentially expressed shoot genes. The discovered motif (CCGGCAGCG) shows significant enrichment in differentially expressed genes relative to non-differentially expressed genes ($P$ = 0.002). For each TFBS, the table reports the TFBS name (motif ID), motif name, TF class, $P$-value, and the number of overlapping base pairs between the discovered motif and the TFBS motif sequence.

| Discovered Motif | TFBS name | Motif name | Class | *P*-value | Overlap (bp) | TFBS motif sequence |
|---|---|---|---|---|---|---|
| CCGGCAGCG | MA1049.1 | ERF094 | AP2/EREBP | 0.000723125 | 8 | CGGCGGCG |
| CCGGCAGCG | MA1034.1 | Os05g0497200 | AP2/EREBP | 0.00143989 | 8 | TGGCGGCG |
| CCGGCAGCG | MA1266.1 | RAP2-11 | AP2/EREBP | 0.00163309 | 9 | CGCCGGCGGCGGCGG |
| CCGGCAGCG | MA1818.1 | Zm00001d052229 | AP2/EREBP | 0.00183892 | 9 | GGCGGCGGCG |
| CCGGCAGCG | MA1006.1 | ERF6 | AP2/EREBP | 0.0024141 | 7 | ACGCCGGCAG |
| CCGGCAGCG | MA1832.1 | Zm00001d002364 | AP2/EREBP | 0.00241669 | 9 | GGCGGCGGCGG |
| CCGGCAGCG | MA1819.1 | Zm00001d005892 | AP2/EREBP | 0.0024797 | 9 | GGCGGCGGCGGC |
| CCGGCAGCG | MA1821.1 | Zm00001d020595 | AP2/EREBP | 0.0026975 | 9 | GGCGGCGGCGG |
| CCGGCAGCG | MA1817.1 | Zm00001d020267 | AP2/EREBP | 0.00320207 | 9 | GCGGCGGCGGCGG |
| CCGGCAGCG | MA1004.1 | ERF13 | AP2/EREBP | 0.0033821 | 8 | TGGCGGCG |
| CCGGCAGCG | MA1671.1 | ERF118 | AP2/EREBP | 0.00351031 | 8 | CGGCGGCGGCG |
| CCGGCAGCG | MA0979.1 | ERF008 | AP2/EREBP | 0.00356266 | 8 | GGGCGGCG |
| CCGGCAGCG | MA0567.1 | ERF1B | AP2/EREBP | 0.0041824 | 8 | TGGCGGCG |
| CCGGCAGCG | MA1820.1 | Zm00001d024324 | AP2/EREBP | 0.00442989 | 9 | CGGCGGCGGCGGC |
| CCGGCAGCG | MA1833.1 | Zm00001d049364 | AP2/EREBP | 0.00493983 | 9 | GGCGGCGGCGGCGGCG |
| CCGGCAGCG | MA1041.1 | MYB55 | Tryptophan cluster factors | 0.00587486 | 8 | CGGTAGGT |
| CCGGCAGCG | MA1005.2 | ERF3 | AP2/EREBP | 0.00619531 | 9 | GCGGCGGCGGC |
| CCGGCAGCG | MA1245.2 | ERF112 | AP2/EREBP | 0.00619531 | 9 | GCGGCGGCGGC |
| CCGGCAGCG | MA0976.2 | CRF4 | AP2/EREBP | 0.00656996 | 8 | CGGTGGCGGCGG |
| CCGGCAGCG | MA1804.1 | WIN1 | AP2/EREBP | 0.0066693 | 9 | ACGGCGGCGG |
| CCGGCAGCG | MA1242.1 | DREB2F | AP2/EREBP | 0.00706261 | 9 | CCGCCACCGCC |
| CCGGCAGCG | MA2013.1 | ERF115 | AP2/EREBP | 0.00736938 | 9 | GCTGCGGCGGCGACG |
| CCGGCAGCG | MA1001.3 | ERF11 | AP2/EREBP | 0.00752188 | 8 | CGGTGGCGGCGG |
| CCGGCAGCG | MA0975.1 | CRF2 | AP2/EREBP | 0.00775011 | 7 | GGCGGCGG |
| CCGGCAGCG | MA2022.1 | LOB | Other C4 zinc finger-type factors | 0.00840717 | 9 | GGCGGCGGCGGCGGCGG |
| CCGGCAGCG | MA1225.1 | ERF5 | AP2/EREBP | 0.00894643 | 9 | TGGCGGCGGCGGTGG |
| CCGGCAGCG | MA1053.1 | ERF109 | AP2/EREBP | 0.00905545 | 7 | GGCGGCGC |
| CCGGCAGCG | MA0997.1 | ERF069 | AP2/EREBP | 0.00932805 | 8 | TGGCGGCGC |
| CCGGCAGCG | MA1264.1 | ERF095 | AP2/EREBP | 0.00937656 | 9 | CGGCGGCGGCGGCGG |
| CCGGCAGCG | MA1233.2 | ERF021 | AP2/EREBP | 0.00982209 | 9 | TTTTGTCGGCGGCGG |
| CCGGCAGCG | MA1051.1 | RAP2-3 | AP2/EREBP | 0.0100017 | 7 | GGCGGCGC |

| | | | | | | |
|---|---|---|---|---|---|---|
| CCGGCAGCG | MA1228.1 | ERF091 | AP2/EREBP | 0.0123693 | 9 | GCGGCGGCGGCGGCGGC |
| CCGGCAGCG | MA0994.2 | ERF8 | AP2/EREBP | 0.0124322 | 9 | CGGTGACGGCGGCGGTGG |
| CCGGCAGCG | MA1000.2 | ERF105 | AP2/EREBP | 0.012844 | 9 | GTGGCGGCGGCGGAG |
| CCGGCAGCG | MA1247.1 | ERF087 | AP2/EREBP | 0.0139954 | 9 | TGGCGGCGGCGGTGG |
| CCGGCAGCG | MA0123.1 | abi4 | AP2/EREBP | 0.0142127 | 9 | GGGGGCACCG |
| CCGGCAGCG | MA1262.1 | ERF2 | AP2/EREBP | 0.0144691 | 9 | TGGCGGCGGCGGAGGCGGCGG |
| CCGGCAGCG | MA2039.1 | ASIL2 | AP2/EREBP | 0.0144695 | 9 | ATCTCCGGCGACG |
| CCGGCAGCG | MA1221.1 | RAP2-6 | AP2/EREBP | 0.015226 | 9 | TGGCGGCGGCGGAGG |
| CCGGCAGCG | MA1252.1 | PUCHI | AP2/EREBP | 0.015226 | 9 | TGGCGGCGGCGGCGG |

**Table S3**

Differentially accumulated metabolites in root and shoot tissue of CRISPR-Cas9 RTx430 Sb*CCD8b* deletion (*ccd8b*) mutants.

| Tissue | ID | Up/Down regulated | m/z | Retention time | *P*-value | Putative annotation |
|---|---|---|---|---|---|---|
| Root | 120 | Down | 479.0846 | 1.006946 | 0.006392 | |
| Root | 392 | Up | 159.9692 | 4.165591 | 0.000571 | DIHYDROOROTATE |
| Root | 1857 | Up | 147.0682 | 19.14538 | 0.000653 | |
| Root | 1789 | Up | 202.1087 | 18.47195 | 0.001547 | |
| Root | 1818 | Up | 137.0822 | 18.56252 | 0.005556 | |
| Root | 1786 | Up | 186.0798 | 18.43877 | 0.005685 | |
| Root | 481 | Up | 131.9743 | 5.295086 | 0.005945 | |
| Root | 511 | Up | 182.9852 | 5.87326 | 0.004698 | |
| Root | 1205 | Up | 113.9637 | 13.05394 | 0.005532 | |
| Root | 1197 | Up | 154.9902 | 13.03924 | 0.004872 | |
| Shoot | 1098 | Down | 161.0822 | 18.78025 | 0.037635852 | |
| Shoot | 412 | Down | 113.9637 | 7.685511 | 0.029374685 | |
| Shoot | 1067 | Down | 163.0978 | 18.52206 | 0.032746445 | |
| Shoot | 1104 | Down | 136.0869 | 18.81891 | 0.037823209 | |
| Shoot | 841 | Down | 163.0978 | 16.07707 | 0.004542073 | |
| Shoot | 817 | Up | 167.0927 | 15.86226 | 0.044182506 | |
| Shoot | 1091 | Up | 120.0556 | 18.74222 | 0.03389208 | |
| Shoot | 172 | Up | 334.0897 | 1.004966 | 0.022774035 | |

**Table S4**

Differentially accumulated metabolites in root and shoot tissue of CRISPR-Cas9 Macia *LGS1* deletion mutants (*lgs1-d*) of the $S_2$ generation from three transformation events (V1,2,3).

| Tissue | ID | Deletion event | Up/Down regulated | m/z | Retention time | *P*-value | Putative annotation |
|---|---|---|---|---|---|---|---|
| Root | 1596 | V1 | Down | 161.0822 | 17.04002 | 7.22E-06 | |
| Root | 1721 | V1 | Down | 152.0818 | 17.90326 | 0.003878544 | |
| Root | 1185 | V1 | Down | 115.0502 | 12.99056 | 0.003768248 | |
| Root | 1840 | V1 | Down | 148.0869 | 18.73577 | 0.00377854 | |
| Root | 372 | V1 | Down | 111.0616 | 3.81812 | 0.005499904 | |
| Root | 1565 | V1 | Down | 110.0713 | 16.74528 | 0.007022313 | |
| Root | 253 | V1 | Down | 134.0706 | 1.758279 | 0.005397558 | |
| Root | 199 | V1 | Down | 141.074 | 1.229471 | 0.013924071 | |
| Root | 1752 | V1 | Down | 109.076 | 18.10745 | 0.013935346 | 2,6-Dimethylpyrazine |
| Root | 628 | V1 | Up | 154.9902 | 7.554572 | 0.013309697 | |
| Root | 1361 | V2 | Down | 109.076 | 14.90119 | 0.00116316 | 2,6-Dimethylpyrazine |
| Root | 1524 | V2 | Down | 111.0807 | 16.51527 | 0.003260371 | |
| Root | 253 | V2 | Down | 134.0706 | 1.758279 | 0.007178619 | |
| Root | 1467 | V2 | Down | 165.1135 | 15.82587 | 0.001700964 | |
| Root | 1355 | V2 | Down | 110.0713 | 14.82448 | 0.007402019 | |
| Root | 165 | V2 | Down | 145.9899 | 1.214996 | 0.001258758 | |
| Root | 1800 | V2 | Down | 107.0604 | 18.49562 | 0.005116554 | |
| Root | 1363 | V2 | Down | 137.0822 | 14.91063 | 0.002731161 | |
| Root | 1803 | V2 | Down | 189.1134 | 18.52012 | 0.0065742 | |
| Root | 1366 | V2 | Down | 186.0848 | 14.97233 | 0.007622491 | |
| Root | 1301 | V3 | Down | 151.1003 | 14.03193 | 0.000574332 | |
| Root | 754 | V3 | Down | 110.0692 | 9.357434 | 0.001439045 | |
| Root | 1361 | V3 | Down | 109.076 | 14.90119 | 0.004672829 | 2,6-Dimethylpyrazine |
| Root | 1364 | V3 | Down | 105.0447 | 15.00413 | 0.005640571 | 3-Cyanopyridine |
| Root | 1715 | V3 | Down | 163.0978 | 17.89829 | 0.004406488 | |
| Root | 1392 | V3 | Down | 149.0606 | 15.14319 | 0.001494871 | |
| Root | 29 | V3 | Up | 137.9875 | 0.424846 | 0.003251323 | |
| Root | 1999 | V3 | Up | 116.9719 | 19.8538 | 0.003444883 | |
| Root | 1959 | V3 | Up | 271.9374 | 19.72944 | 0.009544912 | |
| Root | 1005 | V3 | Up | 137.9875 | 11.96375 | 0.000223791 | |
| Shoot | 711 | V1 | Up | 108.0682 | 13.64615 | 0.0245 | |
| Shoot | 765 | V1 | Up | 163.0978 | 15.00569 | 0.019 | |
| Shoot | 799 | V1 | Up | 120.0556 | 15.52352 | 0.0281 | |
| Shoot | 788 | V1 | Up | 189.1135 | 15.45355 | 0.03 | |
| Shoot | 783 | V1 | Up | 136.0869 | 15.37391 | 0.0314 | |

| | | | | | | | |
|---|---|---|---|---|---|---|---|
| Shoot | 762 | V1 | Up | 166.0975 | 14.94231 | 0.0356 | |
| Shoot | 855 | V1 | Up | 121.0725 | 16.20665 | 0.0386 | |
| Shoot | 842 | V1 | Up | 108.0682 | 16.09317 | 0.0345 | |
| Shoot | 820 | V1 | Up | 179.0927 | 15.80812 | 0.0325 | |
| Shoot | 407 | V1 | Up | 274.274 | 7.544738 | 0.029 | |
| Shoot | 1053 | V2 | Up | 202.1087 | 18.39366 | 0.0059 | |
| Shoot | 847 | V2 | Up | 174.09 | 16.1959 | 0.009 | |
| Shoot | 1033 | V2 | Up | 144.9821 | 18.12091 | 0.0148 | |
| Shoot | 842 | V2 | Up | 108.0682 | 16.09317 | 0.013 | |
| Shoot | 981 | V2 | Up | 144.9821 | 17.53765 | 0.0155 | |
| Shoot | 100 | V2 | Up | 105.1103 | 0.801333 | 0.0268 | |
| Shoot | 950 | V2 | Up | 174.09 | 17.32867 | 0.0281 | |
| Shoot | 177 | V2 | Up | 104.107 | 0.826375 | 0.0288 | gamma-Aminobutyric acid |
| Shoot | 974 | V2 | Up | 160.0857 | 17.4576 | 0.0125 | |
| Shoot | 113 | V2 | Up | 136.0618 | 0.913949 | 0.0147 | Adenine |
| Shoot | 628 | V3 | Up | 146.9803 | 12.55224 | 0.0003 | |
| Shoot | 699 | V3 | Up | 165.1135 | 13.27956 | 0.0286 | |
| Shoot | 718 | V3 | Up | 140.0818 | 13.65803 | 0.0486 | |
| Shoot | 1187 | V3 | Up | 146.9804 | 19.62629 | 0.043 | |
| Shoot | 855 | V3 | Up | 121.0725 | 16.20665 | 0.046 | |
| Shoot | 113 | V3 | Up | 136.0618 | 0.913949 | 0.0462 | Adenine |
| Shoot | 711 | V3 | Up | 108.0682 | 13.64615 | 0.0245 | |
| Shoot | 765 | V3 | Up | 163.0978 | 15.00569 | 0.019 | |

**Table S5**

MZMine3 processing parameters in untargeted metabolomics. Roots and shoots were analyzed separately.

| | **Shoots** | **Roots** |
|---|---|---|
| | **Peak Detection** | |
| Noise Level | 4.0E5 Counts | 5.0E4 Counts |
| | **ADAP algorithm** | |
| Minimum group size | 5 | 5 |
| Group intensity threshold | 4.0E5 | 5.0E4 |
| Minimum highest intensity | 4.0E5 | 5.0E4 |
| Scan-to-Scan accuracy | 0.05 m/z or 10.00 ppm | 0.05 m/z or 10.00 ppm |
| | **ADAP Chromatogram resolution** | |
| Signal/Noise threshold | 7 | 8 |
| Minimum feature height | 100 | 100 |
| Coefficient/area threshold | 110 | 110 |
| Peak duration range | 0.00 - 0.10 min | 0.00 - 0.10 min |
| RT wavelength range | 0.00 - 0.10 min | 0.00 - 0.10 min |
| | **13C isotope filter** | |
| m/z tolerance | 0.050 m/z or 10.00 ppm | 0.050 m/z or 10.00 ppm |
| Retention time tolerance | 0.250 min | 0.250 min |
| Maximum charge | 1 | 1 |
| | **Join aligner** | |
| m/z tolerance | 0.050 m/z or 10.00 ppm | 0.050 m/z or 10.00 ppm |
| Weight for m/z | 50 | 50 |
| Retention time tolerance | 0.250 | 0.250 |
| Weight for RT | 50 | 50 |
| Mobility weight | 1 | 1 |
| | **Gap filling/Peak Finder** | |
| Intensity tolerance | 10% | 10% |
| m/z tolerance | 0.050 m/z or 10 ppm | 0.050 m/z or 10 ppm |
| Retention time tolerance | 0.25 min | 0.25 min |
| Minimum data points | 1 | 1 |

**Table S6**

Primers and probes used to detect *LGS1* gene insertion and marker cassettes excision in transformed RTx430 $T_0$ plants.

| Trait | Primer name | Primer orientation | Primer sequence |
|---|---|---|---|
| mo_CRE | 141940 | forward | AGTCAGGAAGAACCTCATGGACAT |
| | 141941 | reverse | CCAGGTGTGCTCGCTGAAC |
| | 141942 | probe | TCCGCGACAGGCAA |
| NPTII_MOD1 | 201730 | forward | TCCCAGGACAGGACCTTTTG |
| | 201731 | reverse | TGCCATGATGCTGACCTTTTC |
| | 201732 | probe | AGCCATCTGGCACCTG |
| ZM-WUS2(ALT1) | 201673 | forward | AGCAGATCCAGCGCATCAC |
| | 201674 | reverse | TGGAACCAGTAGAAGACGTTCTTG |
| | 201675 | probe | CACGGCAAGATCGA |
| PSB1 | 107417 | forward | TGATTCCGATGACTTCGTAGGTT |
| | 119738 | reverse | GCTAATCGTAAGTGACGCTTGGA |
| | 119743 | probe | TAGCTCAAGCCGCTCG |
| PSA2 | 119739 | forward | CATGAAGCGCTCACGGTTACTAT |
| | 119740 | reverse | TCGTACGCTACTGCCACCAA |
| | 119744 | probe | ACGGTTAGCTTCACGACT |
| LEFTBORDER | 165690 | forward | GATCTCGCGGAGGGTAGCA |
| | 165691 | reverse | CGAGGGAGATGATATTTGATCACA |
| | 165692 | probe | TTGTAACGATGACAGAGCG |
